# Influenza A virus infection perturbs host cell glycosylation

**DOI:** 10.64898/2026.02.02.703422

**Authors:** Kyle L. Macauslane, Cassandra L. Pegg, Joy Seitanidou, Georgina McCallum, Lauren E. Steele, Melanie Wu, Julian DJ Sng, Ellesandra C. Noye, Merrina Anugraham, Daniel Kolarich, Kirsty R. Short, Benjamin L. Schulz

**Affiliations:** School of Chemistry and Molecular Biosciences, The University of Queensland, Brisbane, Australia; .Institute for Biomedicine and Glycomics, Griffith University, Gold Coast, Australia; .Australian Infectious Diseases Research Centre, The University of Queensland, Brisbane, Australia

## Abstract

Glycosylation is critical for viral-host cell interactions in influenza A virus (IAV) infection, but we lack a comprehensive understanding of how IAV infection shapes the host glycoproteome and the implications of these changes. Here, we used a liquid chromatography-tandem mass spectrometry (LC-MS/MS) approach to perform proteomic, glycomic, and glycoproteomic characterisation of the dynamic subcellular responses to an *in vitro* time course infection of human A549 cells with two IAV strains (A/X-31, H3N2; and A/Puerto Rico/8/1934, H1N1). IAV infection resulted in only modest changes to the subcellular proteome, but robust and significant changes to the host secreted and organelle glycome and glycoproteome. Infection with either virus resulted in a widespread reduction in sialic acid across the *N*- and *O*-glyco(proteo)me; increased abundance of oligomannose, paucimannose, and phosphorylated glycans; and shorter hybrid/complex glycans. Reduced sialylation was consistent with desialylation of glycans by viral neuraminidase (NA), but with specific features of the glycan and protein controlling the extent of desialylation. Desialylation was greater when glycans were fucosylated; when the sialic acid was attached via an α2,3 linkage or positioned on the α3 arm; on larger, more complex glycans; and when present on proteins that are more accessible to IAV NA. Subtle but prolonged activation of the unfolded protein response in infection led to a doubling of oligomannose *N*-glycosylation. Glycans were shorter in infection, implicating IAV-induced disruption of Golgi glycoprotein flux as a mechanism that reduces host glycoprotein sialylation and promotes virion release, independent of NA activity. Our data provide important insights into the host glycoproteome during influenza virus infection, furthering our understanding of how influenza NA acts upon host glycans, and how cell stresses in infection perturb key mediators of protein stability and function, cell signalling and immunity.

## Introduction

Influenza A viruses (IAV) remain a considerable challenge in global healthcare, with seasonal epidemics of two antigenic subtypes, H1N1 and H3N2, responsible for substantial morbidity and mortality in humans [1]. The burden of disease has been further heightened by occasional pandemic outbreaks of influenza, with four outbreaks having occurred since the start of the 20^th^ century [2]. Although vaccines and antivirals help combat IAV infection, the virus’s rapid evolution via antigenic drift and shift allow it to persist as a global threat [3]. Novel insights into influenza biology and the host response to viral infection are therefore needed.

Protein glycosylation is a critical process in viral and host cell biology alike; changes in the glycoproteome can impact protein folding, stability, localisation, and activity, and also contribute to changes in cell-cell recognition, signalling, and function [4, 5]. Glycosylation is a post-translational modification of secreted and membrane proteins involving the addition of carbohydrate moieties, or glycans, processed by a biosynthetic network of glycosylation enzymes in the endoplasmic reticulum (ER) and Golgi apparatus [6]. The two predominant forms of protein glycosylation are *N-* and *O-*glycosylation [7, 8]. *N-*glycosylation begins within the ER with the transfer of preassembled oligosaccharides to asparagine residues of nascent proteins, particularly those within the consensus sequon N-X-S/T (X ≠ P) [9, 10]. *N-*glycans are trimmed in the ER to yield oligomannose glycans and may be further modified in the Golgi to yield hybrid and complex glycans [11]. *O*-glycosylation occurs primarily in the Golgi and involves the sequential addition of sugars to serine or threonine residues, resulting in numerous core structures with diverse extensions [8].

Protein glycosylation is of particular importance for IAV. Just as these viruses subvert the machinery of infected host cells for protein biosynthesis, so too do they hijack the glycosylation machinery for the modification of the viral surface glycoproteins haemagglutinin (HA) and neuraminidase (NA) [12, 13]. The HA and NA proteins, the major viral surface antigens, use glycosylation to shield immunodominant epitopes and escape immune recognition [14]. IAVs also use sialylated glycoconjugates as the host receptor for viral entry, with HA recognising *N-*acetylneuraminic acid (NeuAc) bound to galactose in α2,6 linkage for human IAVs, and α2,3 for avian IAVs [15]. In contrast, NA is a sialidase that cleaves sialic acids at the cell surface to initiate budding of progeny virions and to prevent virus aggregation [16].

The non-template derived nature of glycan processing necessitates that the cellular glycoproteome is subject to the availability of glycan substrates, regulation of glycosylation enzymes, and architecture of the secretory protein pathway, and hence the overall health of the organism. It is therefore unsurprising that the glycoproteome is perturbed in a variety of diseased states including cancer, autoimmune and chronic inflammatory diseases [17]. The host glycoproteome is also impacted in a variety of infectious diseases [18]. For example, Hepatitis C virus-infected human hepatoma cells have increased fucosylated, sialylated, and complex *N*-glycans [19]; elevated levels of α2,6 sialylation is a marker of disease severity in the serum and lung tissue of COVID-19 patients [20]; and antibodies from individuals with active tuberculosis have increased core fucosylation and decreased galactosylation and sialylation [21].

Desialylation of host cell surface glycoproteins by NA is necessary for viral release in IAV infection [22], but the range and extent of desialylation across the entire host cell glycoproteome is not yet understood. Here, we used integrated proteomics, glycomics, and glycoproteomics to explore the dynamic host glycoproteome response to IAV infection with H1N1 and H3N2 IAVs representative of the predominant IAV subtypes that circulate in humans.

## Methods

### Cell Culture

Adenocarcinomic human alveolar basal epithelial (A549) cells were used for viral time course infections, and subsequent proteomic, glycoproteomic and glycomic analyses. Madin-Darby canine kidney (MDCK) cells were used for IAV plaque forming assays. All cell lines were cultured in Dulbecco’s Modified Eagle’s Medium (DMEM) supplemented with 10% (*v/v*) foetal-bovine serum (FBS) and 1% (*v/v*) 50 U/mL penicillin-streptomycin (Gibco). All cell lines were incubated at 37 °C with 5% CO_2_. All cell lines were purchased from the ATCC.

### Viral Stocks

The IAV strains used for this work were A/Puerto Rico/8/1934 (PR8, H1N1) and A/X-31 (x31, H3N2). Viral stocks were propagated in the allantoic fluid of embryonated chicken eggs as previously described [23]. The use of embryonated chicken eggs was approved by the University of Queensland Animal Ethics Committee (AE000089). Viral stocks were concentrated in 100 kDa Amicon Ultra MWCO filter units (Millipore) and loaded and washed in PBS. Naïve allantoic fluid was also concentrated in the same manner from eggs mock infected with PBS.

### Viral Infections

Infections were performed in serum-free infection media (DMEM supplemented with 1% penicillin-streptomycin). A549 cells were cultured in T175 flasks until over 90% confluent. Cells were infected at a multiplicity of infection (MOI) of 5 for 1 h at 37 °C, after which the cells underwent four washes with PBS and were cultured in serum-free infection media until sample collection.

### Subcellular Fractionation

Subcellular fractionation of cells was completed in its entirety immediately following harvesting. Samples were maintained at 4 °C unless otherwise specified. Secreted protein fractions were collected and stored at -20 °C. Secreted fractions were thawed and concentrated in 10 kDa MWCO Amicon Ultra filter units (Millipore) at 4000 rcf for 30 min, followed by a buffer exchange into 50 mM HEPES. Cells were collected through gentle scraping in chilled PBS, followed by centrifugation at 250 rcf for 3 min. Cells were resuspended in 1 mL of a hypotonic lysis buffer containing 0.25 M sucrose, 2 mM EDTA, 10 mM HEPES (pH 7.4), 2 mM magnesium acetate tetrahydrate and 1 X cOmplete protease inhibitor cocktail (Roche) and incubated on ice for 5 min. Cells were gently lysed using 70 strokes of a pre-cooled Dounce homogeniser pestle “B” (Sigma-Aldrich). Lysates were treated with 500 U of Benzonase nuclease (Sigma-Aldrich) for 20 min at room temperature, followed by 10 min on ice.

Lysates were centrifuged at 250 rcf for 3 min to pellet denatured nucleic acids and cellular debris. The supernatant then underwent differential centrifugation to pellet subcellular fractions, beginning with a centrifugation at 1400 rcf for 10 min to pellet the nuclear fraction. The supernatant of this fraction underwent subsequent centrifugation at 20,000 rcf for 30 min to pellet the crude organelle / membrane fraction. The remaining supernatant was harvested as the cytosolic fraction. Nuclear and organelle protein fraction pellets were resuspended in 50 µL of 50 mM HEPES. Protein concentrations were calculated using Qubit Protein Quantification Assay Kit (Thermo Fisher Scientific).

### Glycomic Sample Preparation

Organelle and secreted protein fractions were reduced and denatured by adjusting samples to 1% SDS, 10 mM DTT and 50 mM Tris-HCl buffer pH 8.0 and incubating at 95 °C for 10 min. Samples were cooled to room temperature and then alkylated with the addition of acrylamide to a final concentration of 25 mM prior to incubation at 30 °C and shaking at 1500 rpm for 1 h. Excess acrylamide was quenched with DTT to an additional concentration of 5 mM. Proteins were precipitated by the addition of four volumes of 1:1 methanol / acetone (*v/v*) and overnight incubation at -20 °C. Insoluble precipitated protein was pelleted by centrifugation at 18,000 rcf for 10 min, the supernatant was discarded, and the samples were allowed to air dry for 5 min.

Protein pellets were resuspended in 8 M Urea to a concentration of 10 µg/µL. For each sample, 25 µg of protein was blotted onto PVDF membrane (Immobilon-P, 0.22 µm pore, Merck Millipore). *N-* and *O-*glycans were sequentially released as previously described [24]. In brief, *N*-glycans were enzymatically released using PNGase F (NEB, 50 U) incubated at 37 °C for 16 h, and reduced with 2 M NaBH_4_ for 3 hr at 50 °C. From the same PVDF membrane, *O*-glycans were subsequently released and reduced by reductive β-elimination in 1 M NaBH_4_ at 50 °C for 16 h. Reduced *N-* and *O-*glycans were desalted with strong cation exchange chromatography (AG50W-X8 cation exchange resin, BioRad) and carbon cleaned with porous-graphitised carbon (PGC) material (Supelclean EnviCarb, Sigma Aldrich).

### Proteomic Sample Preparation and TMT Labelling

10 µg aliquots of the fractionated protein samples were prepared for tandem mass tag (TMT) labelling as per the manufacturer’s recommended protocol (Thermo Fisher Scientific). In brief, five volumes of lysis buffer (1% SDS in 100 mM triethylammonium bicarbonate (TEAB)) were added to the protein samples, followed by centrifugation at 16,000 rcf for 10 min at 4 °C. The supernatant was transferred to new tubes. Proteins were reduced by the addition of 200 mM tris(2-carboxyethyl)phosphine (TCEP) to a final concentration of 10 mM and incubation at 55 °C for 1 h. Iodoacetamide (IAA) was added to the sample to a final concentration of 17 mM and incubated for 30 min protected from light at room temperature. Proteins were precipitated with the addition of six volumes of chilled acetone and overnight incubation at -20 °C. Precipitated protein was pelleted by centrifugation at 8,000 rcf for 10 min at 4 °C. The supernatant was discarded, and the pellet was allowed to dry for 3 min. Protein pellets were resuspended in 50 mM TEAB for digestion with the addition of 0.25 µg of trypsin and overnight incubation at 37 °C. TMT labelling reagents were equilibrated to room temperature before resuspension in acetonitrile. Digested peptides were labelled at a ratio of 7:1 (TMT reagent : peptide) for 1 h at room temperature. The labelling reaction was quenched by the addition of hydroxylamine to 0.28% and incubation for 15 min at room temperature. Samples were multiplexed and dried using centrifugal evaporation. Proteomic samples were resuspended in 0.1% formic acid (FA) for desalting with C18 ZipTips (Millipore).

### Glycopeptide Enrichment

Glycopeptide enrichment was performed on secreted and organelle protein fractions as previously described [25]. Briefly, 90 µg of multiplexed peptides were resuspended in 0.1% trifluoroacetic acid (TFA) for Sep-Pak C18 (Waters) solid phase extraction. Desalted peptides were eluted in 0.1% TFA in 70% acetonitrile (ACN) and adjusted to 1% TFA in 80% ACN for hydrophilic interaction chromatography (HILIC) glycopeptide enrichment. PolyHYDROXYETHYL A beads (100 Å pore diameter, 12 μm particle diameter; PolyLC) were preconditioned by three washes with ACN, three washes with 1% TFA in water, and three washes with 1% TFA in 80% ACN. The samples were then added to the slurry and incubated for 1 min at room temperature. The slurry was washed three times with 100 μL of 1% TFA in 80% ACN, with the supernatant pooled and collected as the unbound proteomic fraction. The slurry was then washed three times with 100 μL of 1% TFA in water, with the supernatant pooled and collected as the enriched glycopeptide fraction. Samples were dried with vacuum centrifugation and resuspended in 0.1% FA for liquid chromatography-tandem mass spectrometry (LC-MS/MS) analysis.

### Mass Spectrometry

Released glycan mixtures were characterised by PGC-LC-MS/MS essentially as previously described [24]. 4 µL of glycans (equivalent to glycans released from 10 µg protein) were injected on a HyperCarb PGC LC capillary column (1 mm × 30 mm, 3 μm particle size, Thermo) installed on a Dionex UltiMate 3000 UHPLC system coupled with an AmaZon Speed ion trap (Bruker). The solvents used were mobile phase A (10 mM ammonium bicarbonate) and mobile phase B (10 mM ammonium bicarbonate in 70% (v/v) ACN). *N*-glycans were separated over a 65 min gradient as follows: 0-5 min 1% B, 6 min 14% B, 25 min 25% B, 45 min 70% B, 47-50 min 98% B, 51-65 min 1% B at a constant flow rate of 15 µL/min. *O*-glycans were separated over a 65 min gradient as follows: 0-5 min 1% B, 25 min 25% B, 45 min 70% B, 47-50 min 98% B, 51-65 min 1% B. The MS was operated in negative ionisation mode with an ESI spray voltage 3.3 kV, nitrogen gas flow 6 L/min, dry temperature 207°C. MS1 scans were performed in the Ultra Scan mode with scan range of *m/z* 460-1,800, AGC of 7 x 10^4^ ions, 3 micro-scans and a maximum of 200 ms accumulation time. Top 3 precursor ions were selected for MS/MS (scan range *m/z* 100-2,200), AGC 3 x 10^4^ ions, isolation window *m/z* 4, and NCE 35. All MS and MS/MS data were acquired in profile mode and dynamic exclusion was disabled.

Peptides and HILIC enriched glycopeptides were analysed on an Orbitrap Eclipse™Tribrid™ Mass Spectrometer coupled to a Dionex UltiMate 3000 UHPLC. Trapping was performed on an Acclaim PepMap 100 (part number 164750) and gradient elution on an Ion Optics Aurora C18 column (1.7 µm, 250 mm x 75 µm). Peptide and glycopeptide samples were loaded onto the trap and washed for 4 µL/s in 99% solvent A (0.1% FA) and 1% solvent B (0.1% FA in 80% ACN). Peptides and glycopeptides were eluted onto the column and separated with the following gradient over a 120 min total run time at a 0.2 µL/min flow rate: 4% B for 4 min, increased to 8% B at 5 min, ramped to 18% B at 13 min, then to 19% B at 14 min, followed by a linear increase to 28% B at 45 min, 38% B at 65 min, and 60% B at 71 min. The gradient was then ramped to 95% B at 72 min and held until 95% B until 95 min, followed by column re-equilibration at 4% B from 97 min. Survey scans of precursors from *m/z* 600 to 1800 for glycopeptides, and *m/z* 375 to 1800 for peptides were acquired in the Orbitrap at 120K resolution (FWHM) using an automatic gain control target of 1,000,000 and maximum injection time of 75 ms. Glycopeptide precursors from charges 2-8 were selected based on highest charge then lowest *m/z* and fragmented using stepped HCD (25%, 40%, and 55%). Peptide precursors from charges 2-8 were fragmented using stepped HCD (15%, 40%, and 55%). Dynamic exclusion of previously selected ions was enabled for 120 s. Fragment ions were acquired in the Orbitrap at a resolution of 60K using an automatic gain control target of 500,000 and maximum injection time of 250 ms.

### Data Analysis

LC-MS/MS raw glycomics data were manually inspected using Compass Data Analysis v4.0 (Bruker). Monoisotopic precursors were searched against GlycoMod [26] with a mass tolerance of 0.5 Da to identify putative monosaccharide compositions. Only compositions containing hexose (Hex), *N*-acetylhexosamine (HexNAc), deoxyhexose (dHex), sialic acid (NeuAc), and/or phosphorylation were considered. Putative glycan identifications were manually confirmed using MS/MS fragmentation spectra and PGC-LC retention time as previously described [27, 28]. Extracted ion chromatogram (EIC) based, area-under-the-curve glycan quantification was performed using Skyline v.24.1.0.414 [29]. Glycan abundance was normalised as a proportion of all glycans in each sample.

The identification of peptides for proteomic analysis was performed using the Sequest node in Proteome Discoverer (Thermo Fisher Scientific, v2.5.0.400) against a composite database containing all high confidence proteins from the human proteome (downloaded from UniProt 25/11/2021), porcine trypsin, 199 contaminating proteins present in FBS described previously [30], and 15 IAV proteins (13 proteins from A/Puerto Rico/8/1934 as well as the HA and NA surface proteins from A/Aichi/2/1968 downloaded from UniProt 12/04/2022).

Identification of glycopeptides for glycoproteomic analysis was performed using the Byonic (v4.3.4) node of Proteome Discoverer (Thermo Fisher Scientific, v2.5.0.400). Cleavage specificity was set to C-terminal to arginine/lysine permitting 2 missed cleavage events. Mass tolerance was set to 10 ppm for precursor ions and 15 ppm for fragment ions. Fixed modifications were set for carbamidomethylation of cysteines and TMT labelling at peptide N-terminus and lysines. Variable modifications (deamidated asparagine and monooxidised methionine) were set as “Common 1”. A glycomics-informed database (built-in Byonic human 182 no multiple fucose database supplemented with compositions identified in our glycomics analysis) containing 223 human *N*-linked, and 9 common *O*-linked glycans were used for the identification of glycan compositions, with all glycan modifications set as “Rare 1”. The proteomic database described above was also used for the identification of (glyco)peptides in Byonic. Quantification of peptides was performed using the Reporter Ion Quantifier node as previously described [31]. The coisolation threshold was set to 50% for proteomics data analysis and 75% for glycoproteomics. Specific protein normalisation was performed on all human proteins. Statistical comparisons were performed using the R package MSstatsTMT [32]. Gene ontology (GO) term enrichment was performed on all human proteins with significant (Benjamini-Hochberg adjusted p < 0.05) increased or decreased abundance with the universe of all proteins identified in the sample using DAVID [33]. Glycopeptide identification and quantification information were integrated using GlypNirO with a Byonic score cutoff of 200, summarising site-specific normalised glycopeptide abundance [34]. Principal component analysis (PCA) and heatmaps of relative glycoform abundances were generated with ClustVis [35].

### Data Availability

The raw and processed mass spectrometry proteomics and glycoproteomics data are available at ProteomeXchange Consortium via the PRIDE [36] partner repository with the dataset identifier PXD072704. The glycomics mass spectrometry data are available at GlycoPost with the dataset identifier GPST000671.

## Results and Discussion

### IAV infection perturbs the host subcellular proteome

We infected human A549 cells with either the lab-adapted A/Puerto Rico/8/34 H1N1 strain of influenza (PR8), the reassortant A/X31 H3N2 virus containing the H3N2 HA and NA surface glycoprotein genes from A/Aichi/2/1968 and the internal genes of PR8 (x31), or naïve allantoic fluid as a mock infection control. We harvested nuclear, organelle, cytosolic, and secreted protein fractions every 6 h up to 24 h post infection (hpi), and performed TMT-labelled quantitative proteomics on each subcellular fraction at each timepoint to determine the dynamic changes to the proteome in each subcellular compartment during viral infection. Across all time points and subcellular fractions, we identified a total of 3,829 human proteins (Supplementary File 1). Few proteins were significantly altered in abundance in the nuclear or secreted protein fractions at any time point following infection, and early time points in the cytosolic and organelle fractions also demonstrated minimal changes in protein abundance between mock and infected samples (Figure 1A and Supplementary File 1). In contrast, many large and significant changes in protein abundance were observed in the organelle and cytosolic fractions in the later timepoints 12 and 18 hpi (Figure 1A and Supplementary File 1). We performed gene ontology (GO) term enrichment analysis to identify enriched pathways associated with the proteins that changed in abundance in these fractions (Supplementary File 2). Amongst proteins increased in abundance in the cytosol 18 and 24 hpi, we observed enrichment in terms including RNA processing, splicing, and translation (Figure 1B). In the organelle fraction 18 hpi we observed enrichment of similar terms related to mRNA processing in proteins increased in abundance, while the cell adhesion term was enriched in proteins decreased in abundance following infection. These processes have all been previously associated with IAV infection, with the processing of viral RNA and proteins critical to the production of nascent viral particles [37]. There were minimal differences in the proteomes following PR8 (H1N1) or x31 (H3N2) infection (Figure S1 and S2), indicating that the dynamics of the subcellular proteome during infection with these two influenza virus strains were very similar.

**Figure 1.**
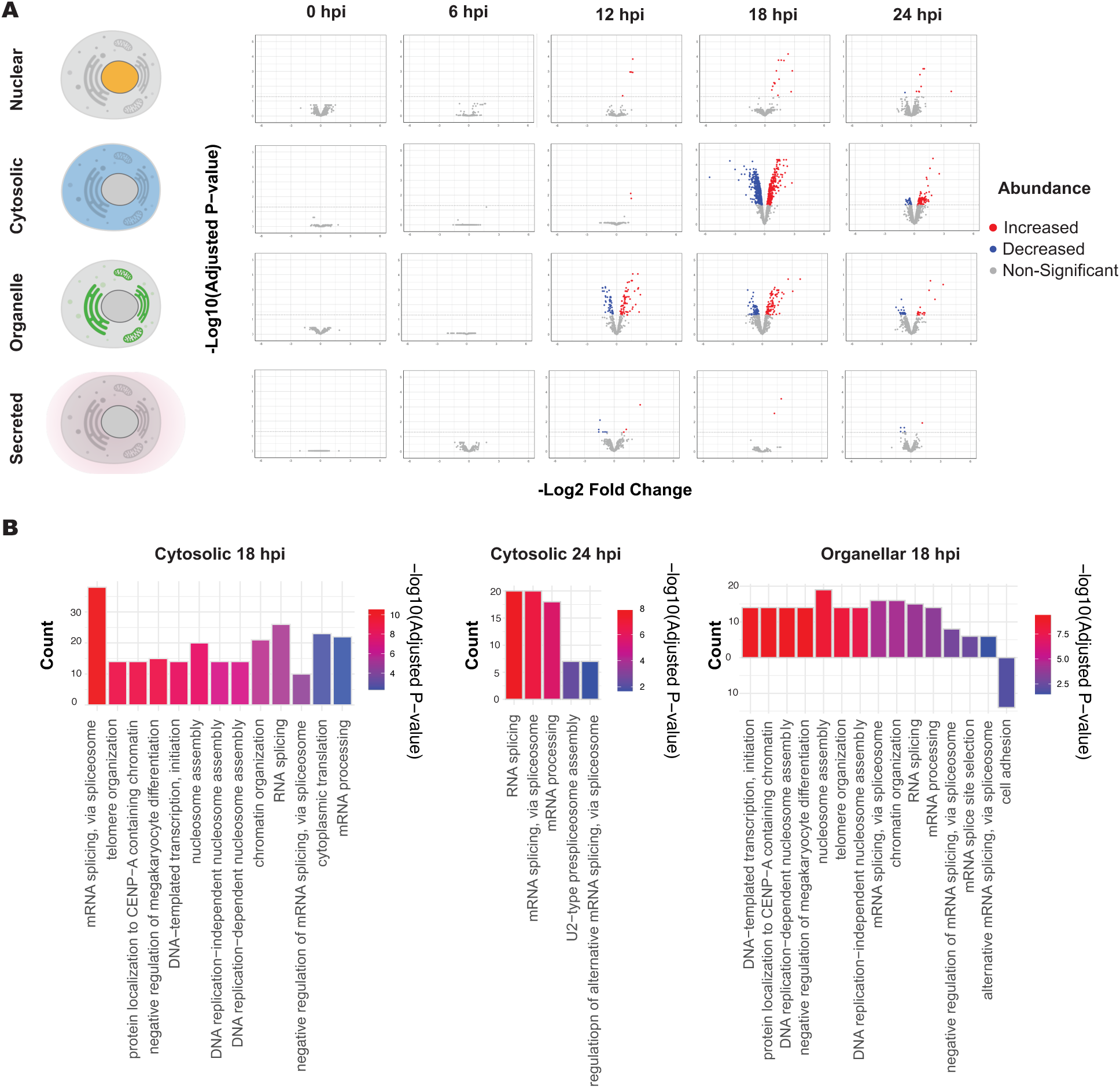
Host-cell subcellular proteomic response to IAV infection. **(A**) Individual volcano plots of differentially abundant proteins are shown for each timepoint (0, 6, 12, 18 and 24 h) post x31 (H3N2) infection in each nuclear, cytosolic, organelle, or secreted fraction. Statistical comparisons were performed using MSstatsTMT. Proteins significantly (adjusted p < 0.05) increased in abundance, red; proteins significantly decreased in abundance, blue; and proteins not significantly different, grey. (**B**) GO Term enrichment of biological processes for significantly increased and decreased proteins following x31 infection. The count of significantly different proteins associated with each term is shown on the y-axis (positive for significantly increased abundance and negative for significantly decreased). Colour gradient indicates the adjusted p-value for enrichment of a term.

### IAV infection strongly perturbs the host cell glycome

As glycosylation is critical during IAV infection, we next considered changes to the host subcellular glycome – that is to say, changes in the abundance, composition and structure of protein-linked glycans. We focused on changes to the *N-* and *O-*glycomes of membrane and secreted proteins following infection. We once again infected A549 cells with the PR8 and x31 viruses, or mock treatment, and at 24 hpi we isolated the organelle and secreted protein fractions; the subcellular fractions expected to contain glycoproteins. *N*-glycans were first released by PNGase F, then *O-*glycans were released by β-elimination. Released *N*- and *O*-glycans were reduced and analysed separately by PGC-LC-MS/MS.

Our analysis of the A549 *N-*glycome identified 142 *N*-glycan structures arising from 70 unique glycan compositions that ranged from low complexity paucimannose and oligomannose glycans to highly processed terminally sialylated complex structures (Supplementary File 3). We performed PCA of the *N*-glycome profiles to assess the variance within the data of mock and infected A549 cells. We observed strong separation between the mock and infected groups, and between the two IAV infections, based on the abundance of organelle (Figure 2A) and secreted *N*-glycans (Figure S3). To determine the key factors driving this separation we stratified the glycan structures, firstly by the major glycan class to which they belonged (paucimannose, oligomannose, hybrid, or complex) and further by their sialylation and fucosylation statuses. A clustered heatmap of the abundance of organelle (Figure 2B) and secreted (Figure S4) *N-*glycans showed two distinct clusters: a cluster of predominantly paucimannose, oligomannose, and neutral complex glycans which were increased in relative abundance in the two infection groups; and a cluster of predominantly sialylated complex glycans which were decreased in infection.

**Figure 2.**
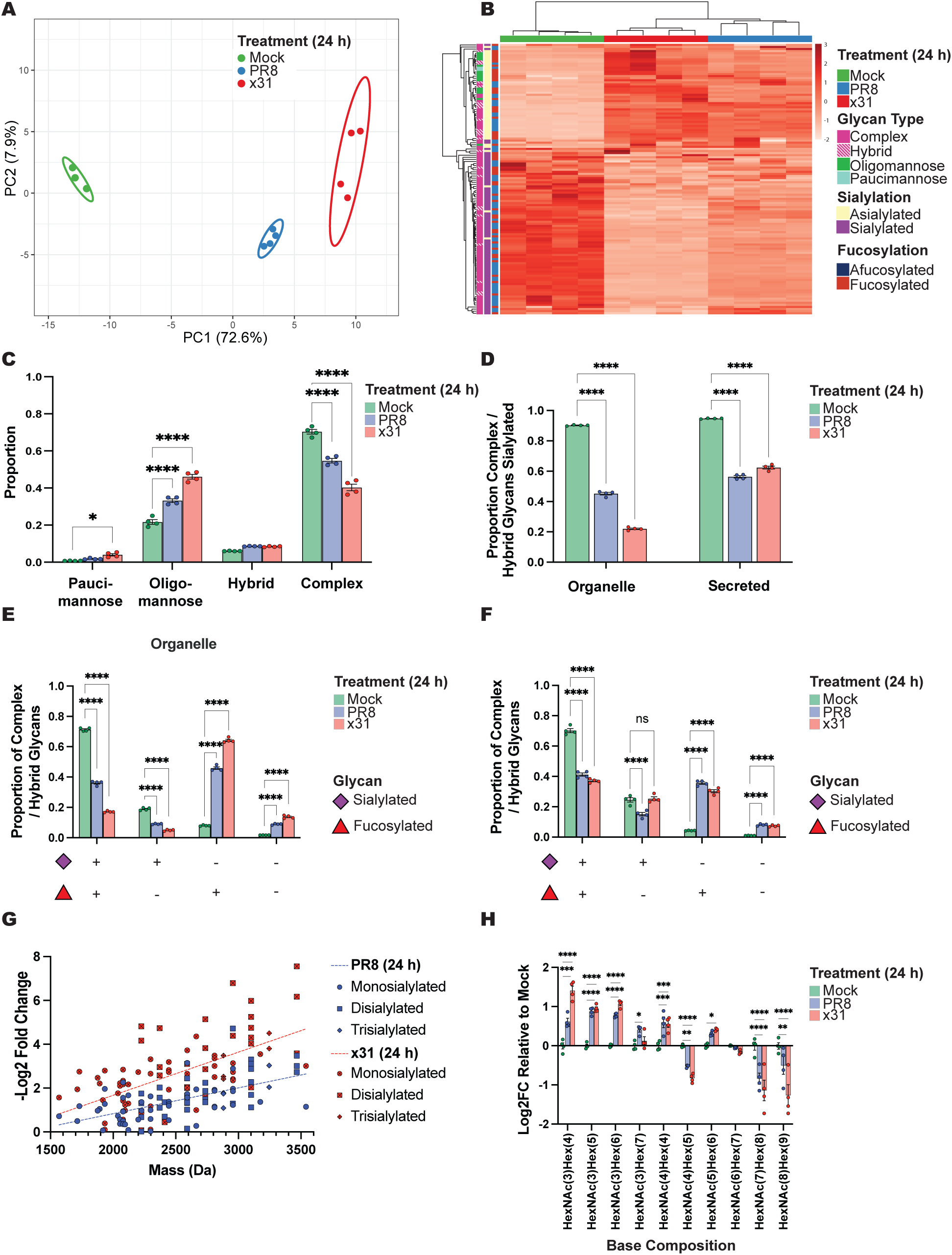
Host-cell *N*-glycome response to IAV infection. **(A)** Principal component analysis of the organelle *N*-glycome of A549 cells 24 hpi with PR8 (H1N1), in blue; x31 (H3N2), in red; and mock infection with naïve allantoic fluid in PBS, in green. *N*-glycans from the organelle *N*-glycome were separated and quantified using PGC-LC-MS/MS and were normalised to total glycan abundance. **(B)** Clustered heatmap of the organelle *N*-glycome, coloured by z-scores of glycan normalised abundance. Glycans are stratified by class, sialylation, and fucosylation status. **(C)** Proportion of each glycan classes in the organelle *N*-glycome. **(D)** Proportion of sialylated complex and hybrid glycans in the organelle and secreted *N*-glycomes. **(E)** Proportion of complex and hybrid glycans in the organelle and **(F)** secreted *N*-glycomes with varying combinations of sialylation and fucosylation. **(G)** Correlation between the decrease in abundance (-Log2 FC) relative to mock of mono-, di- and tri-sialylated organelle *N*-glycans and their mass (Da) following infection. **(H)** Log2FC in abundance relative to mock grouping glycans by HexNAc and Hex compositions (regardless of terminal sialylation or fucosylation). Values in C-F and H show the mean, error bars show SEM. *, P ≤ 0.05; **, P ≤ 0.01; ***, P ≤ 0.001; ****, P ≤ 0.0001.

To investigate potential changes in glycan biosynthesis induced by IAV infection we considered the overall abundance of each major glycan class. Paucimannose and oligomannose *N-*glycans were significantly increased in both PR8 and x31 infections in the organelle (Figure 2C) and secreted fraction (Figure S5), with a greater than 2-fold increase in both glycan classes with x31 in the organelle fraction. This increase is consistent with the increase in cell surface oligomannose glycans that has been observed in severe IAV infections [14]. We also noted that a subset of phosphorylated oligomannose glycans were increased in infection to an even greater extent (Figure S6). Phosphorylated *N*-glycans can target glycoproteins to the lysosome [38], and interestingly can also be used as receptors by IAV [39]. As paucimannose glycans are generated by glycosidases in the lysosome [40], this increase in phosphorylated glycans, and hence lysosomal trafficking, is also consistent with the increase in paucimannose glycans we observed.

The strongest feature driving separation of the glycomes of the infected and mock groups by PCA and clustered heatmap analyses (Figure 2A-B) was the extent of sialylation. However, there were also changes in oligomannose and paucimannose glycans that cannot be sialylated. To specifically examine changes in sialylation independent of changes in the underlying glycan type, we therefore compared the proportion of hybrid and complex type glycans with or without the sialic acid NeuAc. Within the organelle and secreted fractions of mock treated A549 cells, approximately 90% of hybrid and complex glycans were sialylated (Figure 2D). Upon infection with either PR8 or x31 there was a large and significant reduction in this sialylated proportion, dropping to 57.3% / 62.6% for secreted *N*-glycans, and to 47.6% / 27.2% for organelle *N*-glycans for PR8 and x31, respectively (Figure 2D). The strong reduction in the proportion of sialylated glycans in infection is consistent with the enzymatic activity of NA [16]. NA activity is essential for the release of viral progeny at the later stages of the infectious replication cycle as the HA protein would otherwise remain bound to sialylated surface host proteins. It also serves to remove sialylation on HA or NA itself, in turn preventing viral aggregation and promoting efficient propagation of viral progeny. While the function of NA and its role in the replication cycle are well characterised, it is noteworthy that the desialylation we observed in the infected host cell’s organelle and secreted *N*-glycome (Figure 2D) was substantial and widespread, and clearly distinguished infected from uninfected cells.

Although IAV infection induced widespread desialylation of *N*-glycans, not all glycans were equally affected. We therefore investigated the factors that could influence the extent of desialylation of a particular *N*-glycan. We first considered the co-occurrence of fucosylation, the other main terminal glycan modification besides sialylation. To assess this, we once again considered the change in abundance of hybrid and complex glycans, but this time separated into groups based on both sialylation and fucosylation status. We observed that fucosylated glycans were more efficiently desialylated than afucosylated glycans, particularly for secreted *N*-glycans (Figure 2E-F). We next considered the influence of glycan size on the extent of desialylation. For all sialylated organelle and secreted *N-*glycans, we considered the relative fold change in abundance upon infection as a function of the glycans’ mass. We observed a significant correlation between the reduction in relative abundance of these sialylated glycans and glycan mass (significantly non-zero slope, p < 0.0001), with larger *N*-glycans being more efficiently desialylated with IAV infection in the organelle (Figure 2G) and secreted fraction (Figure S7). This is consistent with NA-glycan interactions derived from lectin-based assays that suggest NA interacts best with large and highly branched *N*-glycans [41, 42]. These higher mass glycans are also more likely to contain multiple sialic acids, another factor likely to improve NA interaction and desialylation. However, it was also possible that the efficient desialylation of large glycans was not entirely driven directly by NA desialylation but also by changes in the size of the underlying glycan. To assess this, we stratified hybrid and complex glycans by their size based only on their number of Hex and HexNAc residues, regardless of other monosaccharides. This analysis showed that infection resulted in shorter hybrid/complex *N*-glycans: we observed an increase in the relative abundance of lower mass hybrid structures and a decrease in the abundance of large complex structures in the organelle (Figure 2H) and secreted fraction (Figure S7). These larger structures are more extended, more branched, and require more processing by glycosylation enzymes. That IAV infection results in smaller *N*-glycans is therefore consistent with extended *N*-glycans being particularly susceptible to the various cellular stresses induced by infection. Small *N*-glycans intrinsically have less capacity to be sialylated, suggesting general perturbation of Golgi glycosylation flux is a mechanism by which IAV infection reduces host glycoprotein sialylation to promote virion release, independent of NA activity. In summary, these analyses found that large and fucosylated *N*-glycans were more efficiently desialylated in IAV infection, which also curtailed *N*-glycan extension and branching.

We were curious about the positive effect of fucosylation on the extent of desialylation (Figure 2E-F). To explore this, we therefore next considered the effect that the addition of fucosylation had on the extent of desialylation in defined example glycan structures. We assessed the desialylation of three disialylated bi-antennary glycan structural isomers and identified corresponding structures with core fucosylation (as validated by MS/MS spectral evidence). The core fucosylated structures were indeed more efficiently desialylated than their afucosylated counterparts in IAV infection (Figure 3). Hence, the addition of core fucose, with an otherwise identical glycan composition and structure, directly correlated with an improvement in the extent of desialylation. That the presence of fucose correlated with improved NA activity has not to our knowledge been previously reported. Fucosylation has been reported to impact the interactions of many glycoproteins, such as the reduced efficacy of IgG1 antibodies with core fucosylation to induce antibody dependent cellular cytotoxicity [43]. The exact mechanisms causing fucosylation to enhance interactions with NA here are unknown but likely it causes, or correlates with, increased accessibility of the glycan.

**Figure 3.**
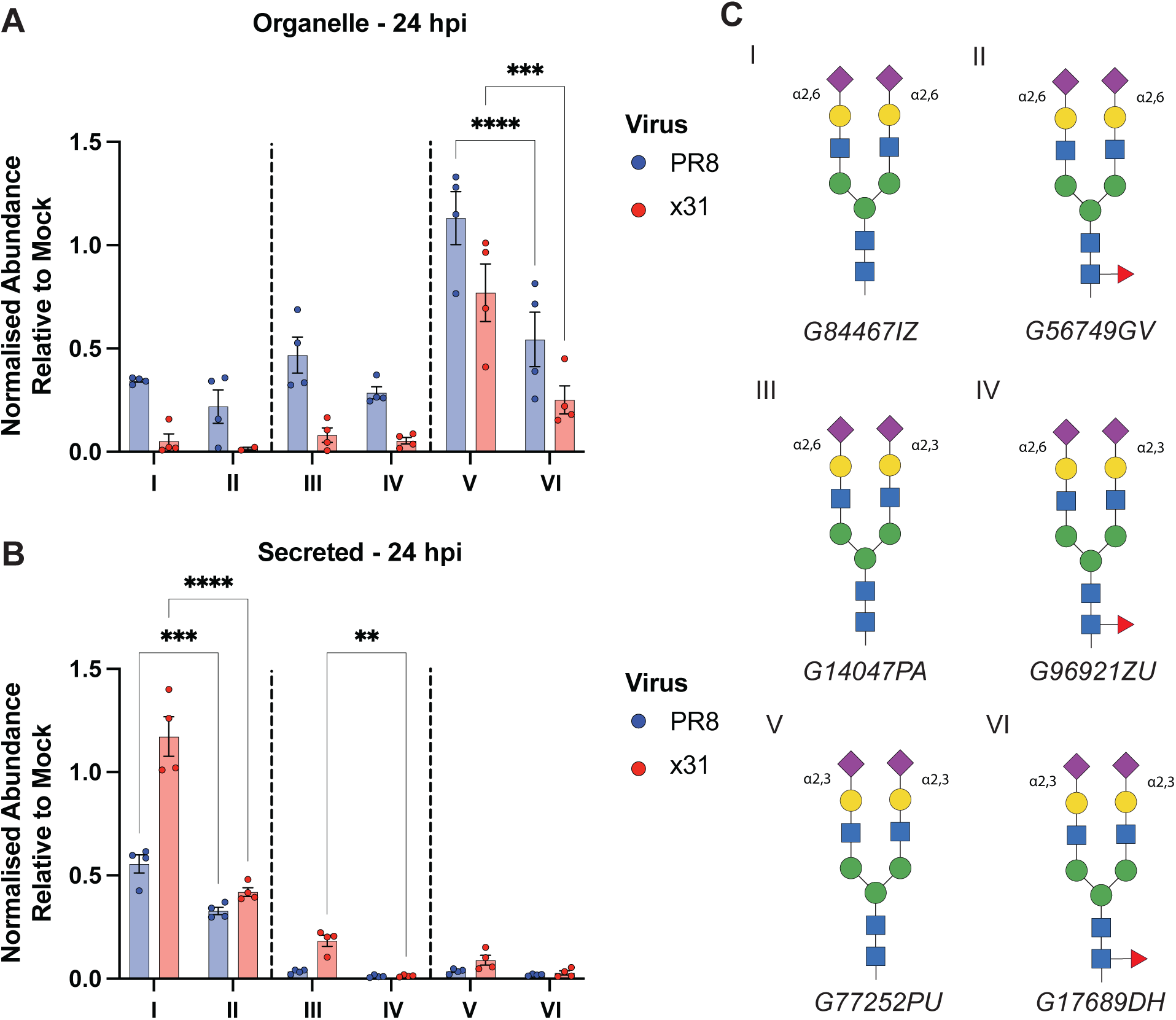
Effect of core fucosylation on the extent of desialylation in IAV infection. **(A)** Abundance of disialylated bi-antennary *N*-glycans with and without core fucosylation 24 hpi with PR8 (H1N1) (blue), or x31 (H3N2) (red) relative to mock infection in organelle and **(B)** secreted fractions. **(C)** Glycan structural assignments determined by PGC-LC retention time and MS/MS fragmentation, with GlyTouCan ID. Glycan isomer abundances are normalised to all complex and hybrid type glycans to account for changes in oligomannose glycan abundances in infection. Values in A and B show the mean, error bars show SEM. *, P ≤ 0.05; **, P ≤ 0.01; ***, P ≤ 0.001; ****, P ≤ 0.0001.

The precise glycosidic linkage of sialic acid is critical for determining the host range, transmissibility, and tissue tropism of IAV. PGC-LC-MS/MS glycomics can characterise these structural features by PGC-LC separation of glycan isomers with α2,3 and α2,6-linked NeuAc [44]. We therefore asked if the extent of desialylation in IAV infection was affected by sialic acid linkage or other glycan structural features. To assess this question, we focussed on a subset of structurally related glycans, di-, mono-, and a-sialylated bi-antennary *N-*glycans. We identified three structural isomers of the monosialylated bi-antennary glycans (Figure 4A), three structural isomers of the disialylated bi-antennary glycans (Figure 4B), and the asialylated bi-antennary glycan

**Figure 4.**
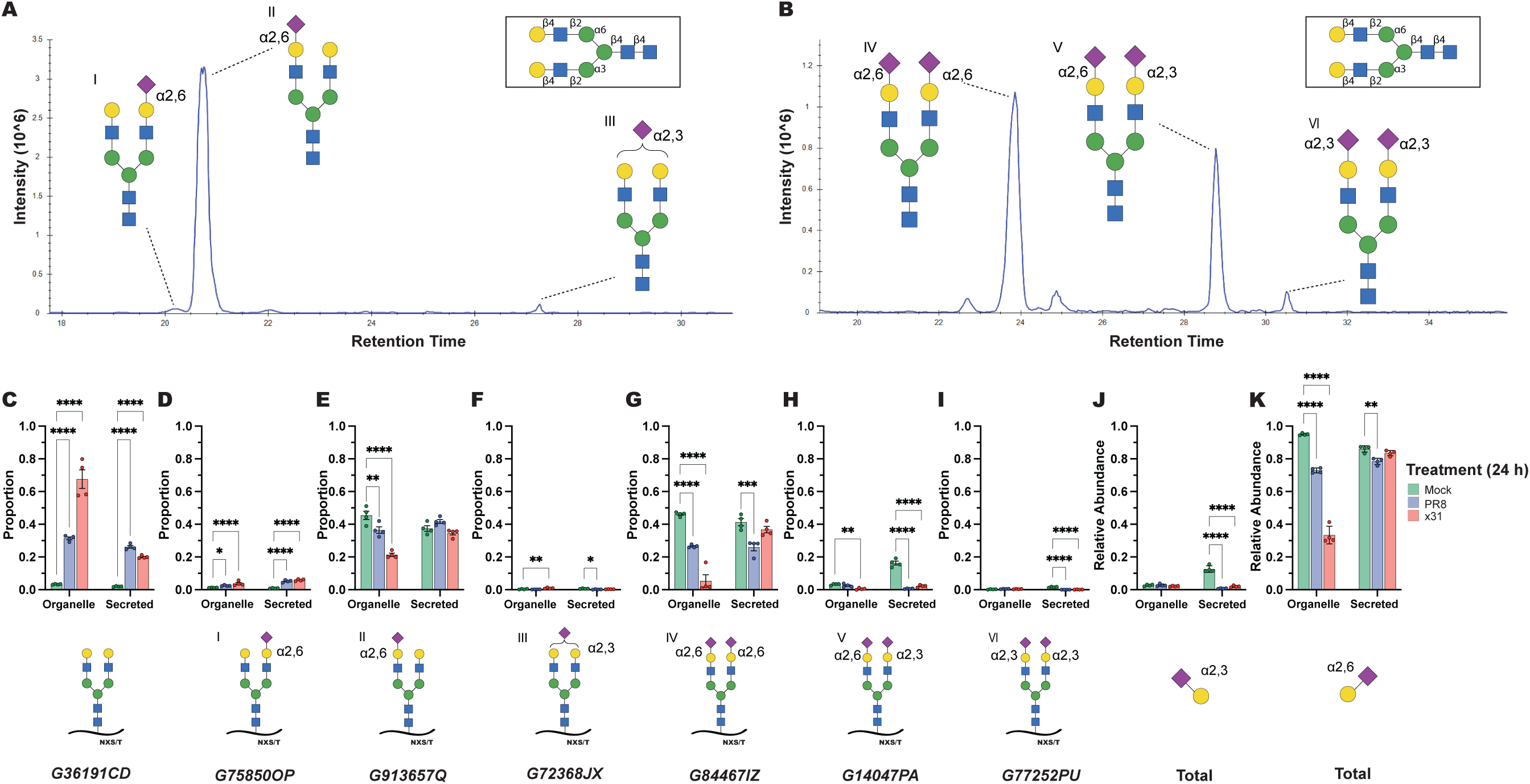
Desialylation of bi-antennary *N*-glycans in IAV infection. **(A)** Extracted ion chromatogram of monosialylated bi-antennary *N*-glycan isomers and **(B)** disialylated bi-antennary *N*-glycan isomers. **(C)** Proportion of the asialyated bi-antennary *N*-glycan 24 hpi infection with PR8 (H1N1) (blue); x31 (H3N2) (red); or mock infection with naïve allantoic fluid (green). **(D-I)** Proportion of the total bi-antennary *N*-glycan abundance for the sialylated glycan structures indicated below each graph. **(J-K)** Relative abundance of *α*2,3 **(J)** or *α*2,6 **(K)** linked NeuAc residues present on the bi-antennary *N*-glycans. Abundance from disialylated glycans was weighted proportionally. GlyTouCan IDs are provided for all unambigous structural assignments. Values in C-K show the mean, error bars show SEM. *, P ≤ 0.05; **, P ≤ 0.01; ***, P ≤ 0.001; ****, P ≤ 0.0001.

We measured the changes in abundance of each of the bi-antennary a-, mono-, and di-sialylated glycan structural isomers after infection. The asialylated glycan increased with both infections in the organelle and secreted fractions (Figure 4C), consistent with it being the complete desialylation product of the larger structures. The behaviour of the monosialylated glycans varied: one structure with α2,6-linked sialic acid on the α6 arm increased in infection (Figure 4D), while the other structures remained steady or decreased (Figure 4E-F). This apparently contradictory behaviour was due to the monosialylated glycans being intermediate products of desialylation of the disialylated glycans. Their abundance was therefore increased by desialylation of disialylated glycans but decreased by their own desialylation. All disialylated structures were decreased in abundance following infection (Figure 4G-I), although the extent of desialylation for each structure varied with each viral strain and/or subcellular location. Bi-antennary glycans with α2,3-linked sialic acid(s) were generally low in abundance initially, particularly in the organelle fraction (Figure 4F and 4I), but were desialylated very efficiently in infection (Figure 4J). Structures with α2,6-linked sialic acid (Figure 4E and 4G) were generally desialylated efficiently in x31 infection in the organelle fraction, but not in the secreted fraction (Figure 4K).

The network of changes to the abundance of these bi-antennary glycan isomers allowed us to dissect how glycan structure shapes desialylation of host glycans by viral NA. Sialic acid on the α3 arm was efficiently desialylated while sialic acid on the α6 arm was protected (Figure 4D-E). This difference is likely due to the shift in conformational equilibrium of the α6 arm from outstretched to folded when it is galactosylated [45]. This folded conformation renders the α6 arm less accessible to sialyltransferases [45] and, our data indicate, to the viral NA as well. We observed that NA cleaved both α2,3- and α2,6-linked sialic acid, but with a clear preference for the α2,3-linkage (Figure 4J-K). HA from human IAVs preferentially bind sialic acid α2,6-linked to galactose, whereas HA from avian IAVs instead prefer α2,3-linked sialic acids [15]. NA proteins are less selective, and are able to cleave both linkages [46], but always with a preference for α2,3-linked sialic acid, consistent with our data (Figure 4J-K). We also noticed a difference in desialylation between x31 and PR8 infection, with x31 infection causing greater loss of α2,6-linked sialic acid (Figure 4K). This is consistent with the receptor preference of human N2 Nas [47] like x31 that are generally well adapted to human (α2,6) receptors. Although the H1N1 PR8 virus is a human IAV, it has been adapted to mice and reacquired preference for α2,3-linked sialic acid [48]. Finally, we observed that x31 infection caused strong desialylation of α2,6-linked sialic acid in the organelle fraction, but no significant change in the secreted fraction (Figure 4K). This is likely driven by differences in host glycoprotein presentation in the secretory pathway and extracellularly, with proteins in the Golgi able to be more efficiently desialylated because they are present at much higher concentration and without the large-scale structural organisation of the extracellular matrix. It is also possible that NA behaves differently in these environments, as Golgi NA is tetrameric, while extracellular NA is primarily incorporated into virions. In summary, IAV infection greatly perturbed the *N-*glycome of the host cells, resulting in increased oligomannose, paucimannose and phosphorylated glycans; decreased branching and length of complex glycans; and desialylation of hybrid and complex glycans governed by glycan accessibility and NA receptor/substrate preference.

We next explored the impact of IAV infection on the *O-*glycome of infected A549 cells. We observed many large and significant changes with IAV infection (Figure 5). We identified 49 *O*-glycan structures from 24 unique glycan compositions (Supplementary File 3). PCA of the organelle *O*-glycome showed that as with the *N*-glycome, the two infected groups strongly separated from mock-infected samples (Figure 5A). Stratification of *O*-glycans by their sialylation and fucosylation statuses in clustered heatmap analyses showed that this separation was overwhelmingly driven by a loss of sialylation with IAV infection (Figure 5B). Interestingly however, a few sialylated *O-*glycans remained unchanged or even increased in relative abundance in infection. Structural characterisation through MS/MS fragmentation and PGC-LC retention time revealed that these *O*-glycans were sialyl-Tn and sialyl-T antigens – small structures with sialic acid linked via α2,6 to the core *N*-acetylgalactosamine (Figure 5C-E). In contrast, *O*-glycans of similar size with a sialic acid in α2,3 linkage to galactose were efficiently desialylated (Figure 5F-G). Together, this suggested that sialic acid linkage was a key determinant of NA substrate specificity of *O*-glycans. Interestingly, as well as being poor substrates for NA from human IAV, these sialyl-Tn and sialyl-T glycan structures are also poorly bound by HA from human IAV [49], while HA from avian IAVs do bind these structures. However, these antigens are tumour-associated and likely to be less abundant in healthy individuals than in A549 cells [50]. Together, our results are consistent with the glycan binding specificity of HA and the sialic acid degrading specificity of NA coevolving in IAV to provide appropriate adhesion and release through the distinct airway glycomes of avian species and humans.

**Figure 5.**
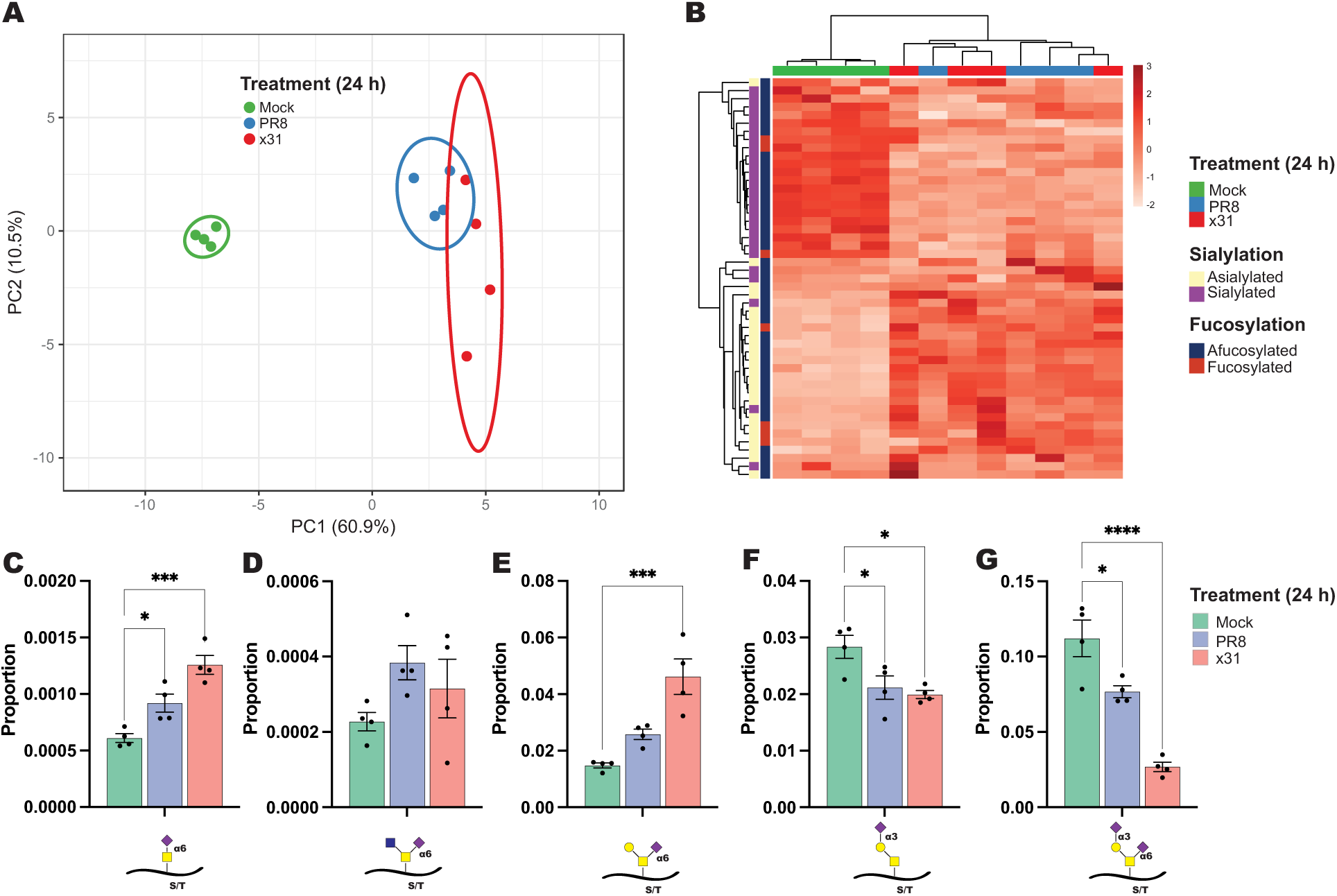
Host-cell *O*-glycome response to IAV infection. **(A)** Principal component analysis of the organelle *O*-glycome of A549 cells 24 hpi with PR8 (H1N1), in blue; x31 (H3N2), in red; and mock infection with naïve allantoic fluid in PBS, in green. *O*-glycans from the organelle *O*-glycome were separated and quantified using PGC-LC-MS/MS and were normalised to total glycan abundance. **(B)** Clustered heatmap of the organelle *O*-glycome coloured by z-scores of glycan normalised abundance. Released glycan structures are stratified by their sialylation and fucosylation status. **(C-G)** Normalised abundance of a series of small, truncated *O*-glycans. Glycan structures were determined by PGC-LC retention time and MS/MS fragmentation and are indicated below the relevant figures. Values in C-G show the mean, error bars show SEM. *, P ≤ 0.05; **, P ≤ 0.01; ***, P ≤ 0.001; ****, P ≤ 0.0001.

We observed several structural features in *N-* and *O-*glycans that enhanced the extent of desialylation by the influenza NA. The position and linkage of the sialic acid, and the presence of core fucosylation all influenced a glycan’s NA substrate activity. Some of these factors that influenced glycan accessibility to NA may be generalisable for all interactors of sialic acid, emphasising the fact that not only was there extensive desialylation of host cell glycoproteins upon IAV infection, but that the remaining sialylated glycoconjugates also likely reflect those most poorly capable of interactions. Differences in our data with PR8 and x31 infection suggest that NA substrate preferences in these viruses closely reflect that of their HA proteins, emphasising that maintaining balance between the two glycoproteins is critical in influenza virology. Strain-dependent changes in protein glycosylation suggest expanding these investigations further to additional strains of concern is warranted. Notably, in addition to desialylation, other changes in glycan processing (class distribution, branching and phosphorylation) were also present following IAV infection.

### IAV infection changes the host glycoproteome

Having determined that IAV infection induced substantial changes to the host cell glycome 24 hpi, we sought to determine whether such changes reflected differences in glycoprotein microheterogeneity, and how these changes developed over the course of an infection. To investigate changes in site-specific glycosylation, we analysed enriched glycopeptides from our TMT labelled secreted and organelle peptide samples from our time-course infections, analysed these with LC-MS/MS, and searched the resulting data to determine the site-normalised abundance of *N-*glycopeptides. Importantly, this approach allowed us to measure changes in site-specific glycosylation independent of any changes to glycoprotein abundance. Overall, we identified 3139 unique glycopeptides across all time points and fractions, including 2117 glycopeptides in organelle fractions, and 1880 in the secreted fraction. These unique glycopeptides comprised 741 different glycosites on a total of 395 glycoproteins (Supplementary File 4).

Consistent with the changes we observed in the glycome and proteome, we observed large and significant changes across the glycoproteome, with a clear increase in the number of glycoforms with significantly different site-normalised abundance and the magnitude of the fold-change of these differences over the course of infection with x31 (Figure 6A) and PR8 (Figure S8). We first asked if the changes we had observed in the glycome in IAV infection could be validated in the glycoproteome. To provide an overview of the changes in the glycoproteome we created a clustered heatmap of glycopeptides and their site-specific abundance (Figure 6B). At 24 hpi in the organelle fraction, and consistent with the glycomics data, two distinct clusters drove separation of mock and infected samples: one cluster of predominately sialylated glycopeptides (54.6% sialylated) was decreased in infection, while another cluster featuring mostly asialylated glycopeptides (12.6% sialylated) was increased in infection (Figure 6B). Similar separation was observed in all time points beyond 12 hpi in both organelle and secreted fractions (Figure S9-S13). To more precisely understand the changes in site-specific glycosylation, we considered sites with the largest LogFC in glycosylation. At these sites we observed that infection caused marked decreases in sialylated glycans, with concomitant increases in the underlying asialylated glycans (Figure 6C-E). In agreement with our glycomics analysis, these changes are consistent with the major impact of IAV infection on host glycosylation being direct desialylation of glycans by NA.

**Figure 6.**
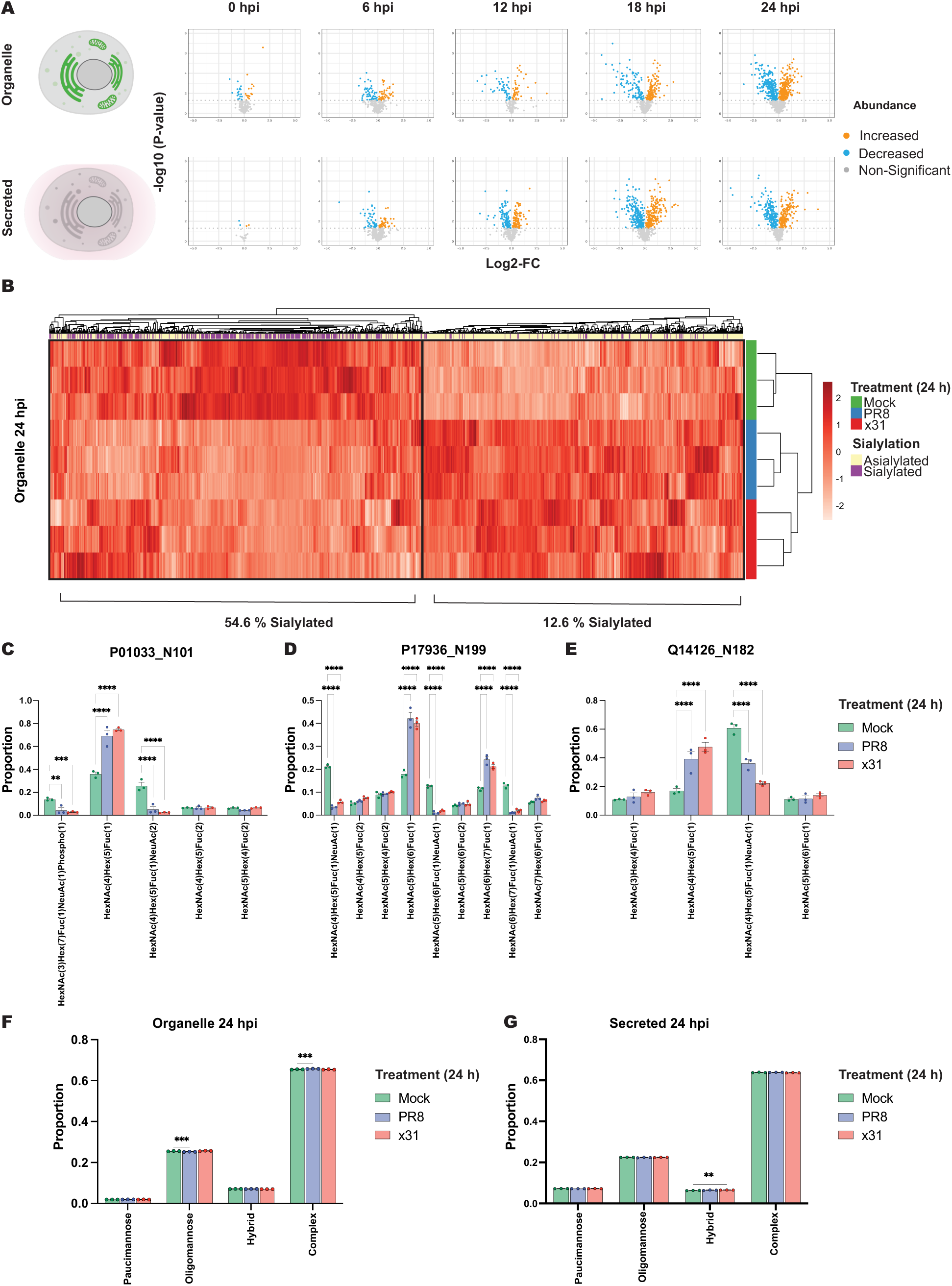
Host-cell *N*-glycoproteome response to IAV infection. **(A)** Volcano plots comparing Log2 FC and p-value of differentially abundant *N*-glycopeptides in organelle and secreted protein fractions every 6 h from 0 to 24 hpi with x31 (H3N2) infection. *N*-glycopeptide abundances are normalised to their site-specific proportions. **(B)** Clustered heatmap of the normalised abundance of organelle *N*-glycopeptides stratified by their sialylation status 24 hpi with PR8 (H1N1), in blue; x31 (H3N2), in red; and mock infection with naïve allantoic fluid in PBS, in green analysed using ClustVis coloured by z-scores of glycopeptide site-normalised abundance. **(C-E)** Glycosylation profiles 24 hpi of selected glycosylation sites: **(C)** N101 of Metalloproteinase inhibitor 1 (P01033), **(D)** N199 of Insulin-like growth factor-binding protein 3 (P17936) and **(E)** N182 of Desmoglein-2. Site-normalised abundance of all identified glycoforms for that site are shown. **(F-G)** Average proportion of major *N*-glycan classes at identified *N*-glycosylation sites 24 hpi in organelle **(F)** and secreted **(G)** protein fractions. Values in C-G show the mean, error bars show SEM. *, P ≤ 0.05; **, P ≤ 0.01; ***, P ≤ 0.001; ****, P ≤ 0.0001.

We next asked if the shift towards oligomannose *N*-glycans in the glycome (Figure 2C) was reflected in site-specific glycosylation. To do this, we stratified glycopeptides by their *N-*glycan classes and compared changes in their summed abundance. While we had observed an increase in oligomannose *N*-glycans in the glycome (Figure 2C), analysis of the glycoproteome showed that site-specific proportions of the various glycan classes remained very consistent across all treatments in both organelle and secreted fractions (Figure 6F-G). This apparent discrepancy between glycomic and glycoproteomic results is consistent with the site-specific glycosylation of these proteins being unaffected by IAV infection, but rather glycoproteins with oligomannose glycans substantially increasing in abundance in IAV infection and driving the global change in the glycome. This increased abundance of oligomannose-modified glycoproteins may be due to newly synthesised viral surface glycoproteins HA and NA, that are glycosylated with some oligomannose *N*-glycans [51, 52]. However, it is unlikely that the viral glycoproteins could alone contribute to the overwhelming increase in oligomannose glycans we observed in the glycome. Indeed, the relative abundance of the HA protein never exceeded 1% of the organelle proteome abundance (Figure S14), whereas the overall proportion of oligomannose glycans doubled in infection to over 40%. Instead, infection with IAV and indeed other respiratory viruses has been reported to increase oligomannose *N*-glycan abundance through activation of the unfolded protein response (UPR), subsequent expansion of the ER [53, 54], and hence increased abundance of ER resident oligomannose-modified glycoproteins. As our proteomic analyses was performed after subcellular fractionation it was not optimal for detecting expansion of the ER, which dominated the organelle fraction. Although the ER was not an enriched cellular compartment in the organelle proteome in infection according to GO term enrichment analysis, plasma membrane (GO:0005886) was enriched in proteins significantly decreased in abundance in infection (Supplementary File 2), consistent with an expansion of the ER at its expense. To assess the extent of UPR activation in our model of IAV infection, we assayed the abundance and expression of UPR-associated proteins in our proteomic analysis, and UPR-associated genes using quantitative reverse translation polymerase chain reaction (qRT-PCR) and identified small but significant UPR activation in IAV infection, consistent with this model (Figure S15-16). Indeed, it is striking that we observed over double the proportion of oligomannose glycans in IAV infection, even with such subtle activation of the UPR, indicating that small but continued UPR activation can result in substantial remodelling of the ER.

### Glycan and protein features control the dynamics of desialylation

Our glycomics analyses established that sialylated glycans were desialylated variably depending on specific glycan features (Figure 2-5); we next validated these effects using glycoproteomics, which also extended our analyses over the course of infection. We focused on the factors that correlated with efficient desialylation in our glycomics data: fucosylation (Figure 2E-F and 3) and glycan size (Figure 2G). We measured the fold change in site-specific abundance of all sialylated glycopeptides at each timepoint and first compared fucosylated vs afucosylated glycans. Consistent with the glycomics data, the presence of fucose significantly increased desialylation, with this effect observed most strongly at later timepoints 18 and 24 hpi (Figure 7A), which were also the time points with the largest overall decrease in sialylation (Figure 7B). We also confirmed that the impact of fucosylation on desialylation was not an artefact of altered fucosylation during infection, since overall fucosylation remained stable throughout the entire course of infection (Figure 7C). We therefore confirmed that the presence of fucosylation enhanced desialylation of *N*-glycans during IAV infection.

**Figure 7.**
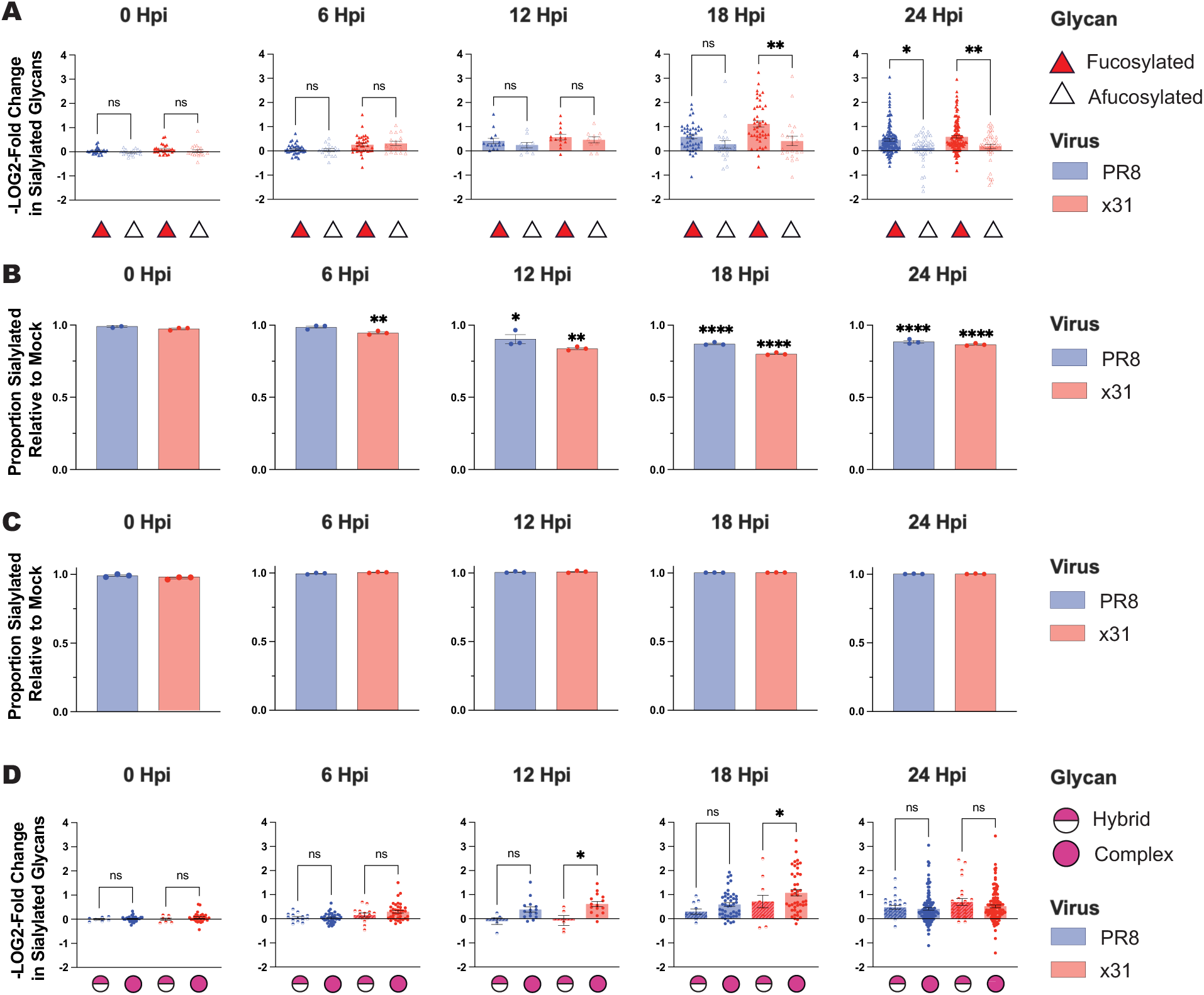
Glycan features controlling desialylation. **(A)** -Log2FC in the site-normalised abundance of sialylated glycopeptides with and without fucosylation in the organelle fraction following a time course IAV infection with PR8 (H1N1), in blue; and x31 (H3N2), in red; relative to mock infection with naïve allantoic fluid in PBS. **(B)** Proportion relative to the average value of mock infected samples of hybrid and complex glycopeptides modified with the sialic acid (NeuAc). **(C)** Proportion relative to the average value of mock infected samples of hybrid and complex glycopeptides modified with fucose. **(D)** -Log2FC in the site normalised abundance relative to mock infection of sialylated glycopeptides in hybrid vs complex *N*-glycans. Values in A-D show the mean, error bars show SEM. *, P ≤ 0.05; **, P ≤ 0.01; ***, P ≤ 0.001; ****, P ≤ 0.0001; ns, not significant.

Our glycomics analysis found that large *N*-glycans were more efficiently desialylated during infection (Figure 2G), an effect which we validated with glycoproteomics by comparing the site-specific desialylation of hybrid (smaller) and complex (larger) sialylated glycans. This analysis confirmed that complex glycans were more efficiently desialylated than hybrid glycans early in infection at 12 and 18 hpi (Figure 7D). This is consistent with hybrid glycans being more likely to be located at less accessible sites with reduced interactions with both glycosylation processing enzymes and NA. At 24 hpi there was no longer a difference in desialylation between hybrid and complex glycans, suggesting that although the larger complex glycans are more readily desialylated by NA, given enough time hybrid glycans can also be desialylated just as completely.

Having established that glycan features contributed to the efficiency of desialylation during infection, we next investigated whether the underlying glycosylated protein impacted desialylation. We only considered glycoproteins with multiple identified glycosylation sites and asked if there was a correlation in the extent of desialylation at sites on the same protein. Indeed, we observed that glycans at different sites on the same protein tended to be desialylated to similar extents (Figure 8), and that some glycoproteins were more efficiently desialylated than others (ANOVA analysis indicated a significant difference in the extent of desialylation due to the protein factor (p < 0.0001)). Glycans on the two gel-forming mucins, Mucin-5B (Q9HC84) and Mucin-5AC (P98088), were very poorly desialylated, as were proteoglycans such as Agrin (O00468) and Macrophage colony stimulating factor-1 (P09603), and extracellular matrix proteins such as Fibronectin (P02751) and Laminin subunits (P11047, O15230). The inefficient desialylation of these extracellular multi-protein complexes is consistent with our glycomics analyses that showed that desialylation was strong in the organelle glycome but inefficient in the secreted fraction (Figure 2D). Inefficient desialylation of these proteins that form very large protein complexes or networks is consistent with IAV virions being physically excluded from these extracellular multi-protein networks. Inefficient desialylation of mucin *N*-glycans could also be due to their extensive *O*-glycosylation, which may serve as better or more abundant substrates for viral NA, or because mucin sialic acids are extensively *O-*acetylated, a feature that would make them poor NA substrates [46, 55] but effective as decoy receptors for HA that prevent IAV infection.

**Figure 8.**
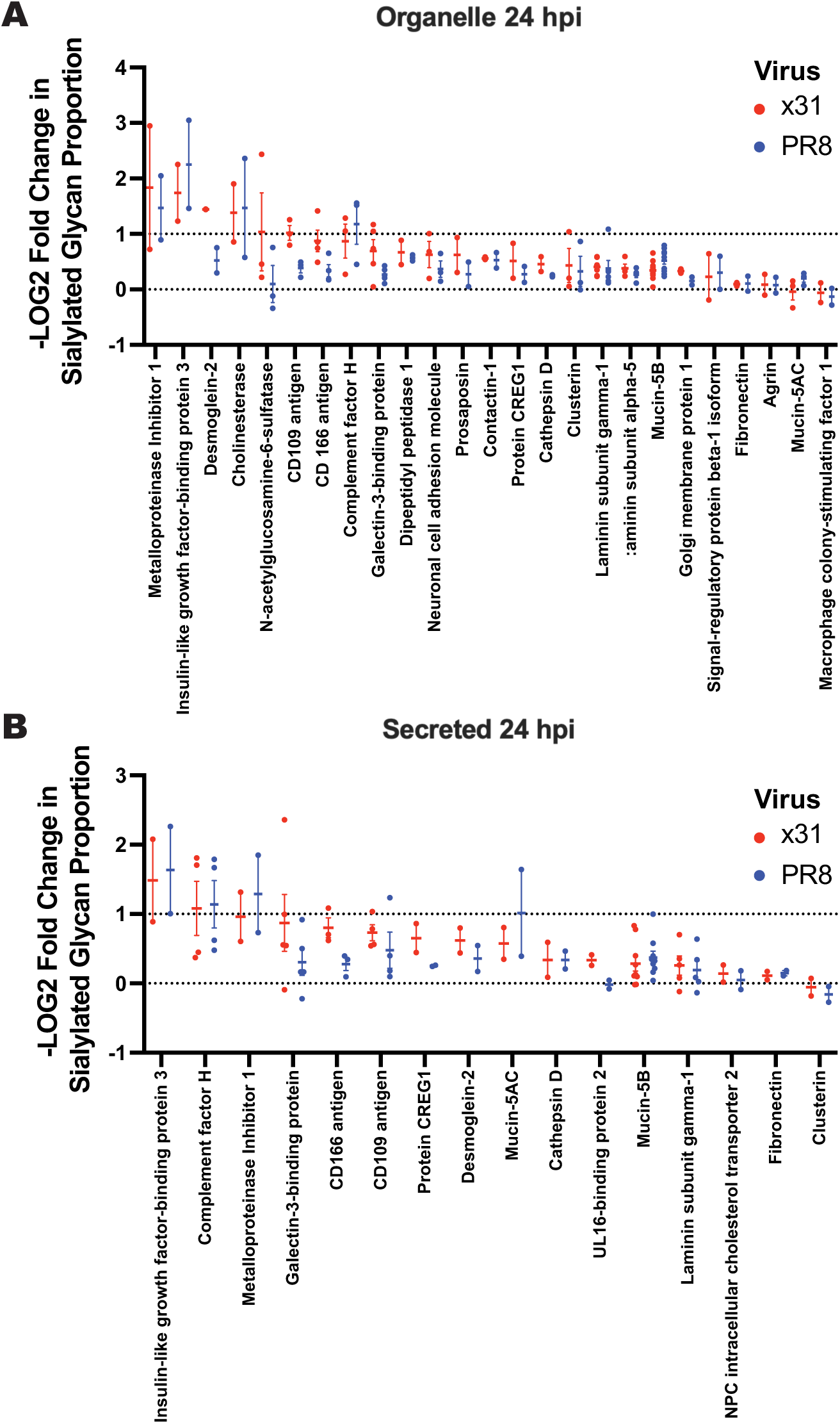
Desialylation of proteins with multiple glycosylation sites. **(A)** -Log2FC in the site normalised abundance of sialylated glycans in organelle and secreted **(B)** protein fractions 24 hpi with x31 (H3N2), in red; and PR8 (H1N1), in blue; relative to mock infection across multiple *N*-glycosylation sites on the same protein where at least 2 sites with sialylated glycans were identified. Values in A-B show the mean, error bars show SEM.

Glycosylation, and sialylation in particular, is critical in regulating the activity of many host proteins, and the substantial desialylation we observe as a result of IAV infection (Figure 2, 5, and 6) is therefore likely to impact their function in diverse and complex ways. For example, Complement Factor H serves as a negative regulator of the complement alternative pathway and as a self-recognition factor by discriminating between “self” cells that display sialic acid and “non-self” bacteria that do not [56]. Siglecs (sialic acid-binding immunoglobulin-like lectins) are primarily expressed on immune cells where they mediate cell-cell interactions [57]. Galectins (galactose binding lectins) are also key regulators of signalling, the immune response, and inflammation, and removal of terminal sialic acid to reveal underlying galactose residues would broadly enhance their activity [58]. Indeed, enhanced galectin-mediated pneumococcal adhesion following IAV infection has been proposed as a contributor to the observed hyper-susceptibility to pneumonia in influenza patients [59]. A decrease in sialylation of proteins secreted from infected cells, including cytokines and other immune signals, could also increase recognition by the hepatic asialoglycoprotein receptor and hence negatively impact their half-life in circulation [60]. These mechanisms are examples that underscore the potential consequences of the extensive global desialylation we observe in organelle and secreted glycoproteins following IAV infection, demonstrating the need for further investigation of the consequences of the dynamic glycoproteome in influenza pathogenesis.

## Conclusion

In summary, we describe the dynamic subcellular proteome, glycome, and glycoproteome of A549 cells following infection with influenza A virus. In addition to general changes to the proteome associated with viral infection, we observed that the host glycome and glycoproteome underwent wide-ranging, large, and significant modifications. We found that there was a prominent global increase in oligomannose glycans due to activation of the unfolded protein response and subsequent expansion of the ER. Further, decreased extension and branching of complex glycans indicated an IAV-driven disruption of Golgi glycosylation flux that presents a mechanism through which IAVs decrease host glycoprotein sialylation to enhance virion release, independently of NA. The largest change in host glycosylation was the substantial widespread desialylation of glycans by viral NA. However, not all glycans were equally desialylated. Several glycan structural features enhanced desialylation through a combination of NA substrate preferences and general glycan accessibility. The presence of fucosylation, position and linkage of the sialic acid, and size of the glycan were all factors that enhanced desialylation of a glycan over the course of infection. The decrease in abundance of certain sialic acid linkages varied between H1N1 (PR8) and H3N2 (x31) infections, in line with the receptor preferences of these viruses, and emphasising that the dynamic glycoproteome in infection was strain-dependent. Another critical factor that influenced desialylation was glycoprotein location, with glycans on large extracellular multi-protein complexes protected from desialylation compared to those on intracellular glycoproteins, likely due to physical shielding and inaccessibility to viral NA. The degree of desialylation we observe, and the factors we demonstrate to be important in its control, reframe our understanding of the impact of NA activity, and IAV infection broadly, on the infected host cell. IAV infection profoundly perturbs the glycosylation of infected host cells, and understanding the functional consequences of these changes in glycosylation will provide critical insights into mechanisms of influenza virology and pathogenesis.

## Supporting information

Supplementary File 1 Proteomics Raw

Supplementary File 2 GO Term Analysis

Supplementary File 3 Glycomics

Supplementary File 4 Glycoproteomics GlypNiro

## Acknowledgements

We thank The University of Queensland, School of Chemistry and Molecular Biosciences Mass Spectrometry Facility for assistance and expertise. We acknowledge the Queensland Node of Metabolomics and Proteomics Australia for their assistance with proteomics data acquisition. This work was supported by a National Health and Medical Research Council (NHMRC) Ideas grant APP1186699 to BLS, CLP, and KRS.

