## Supplementary Figures for "Influenza A virus infection perturbs host cell glycosylation"

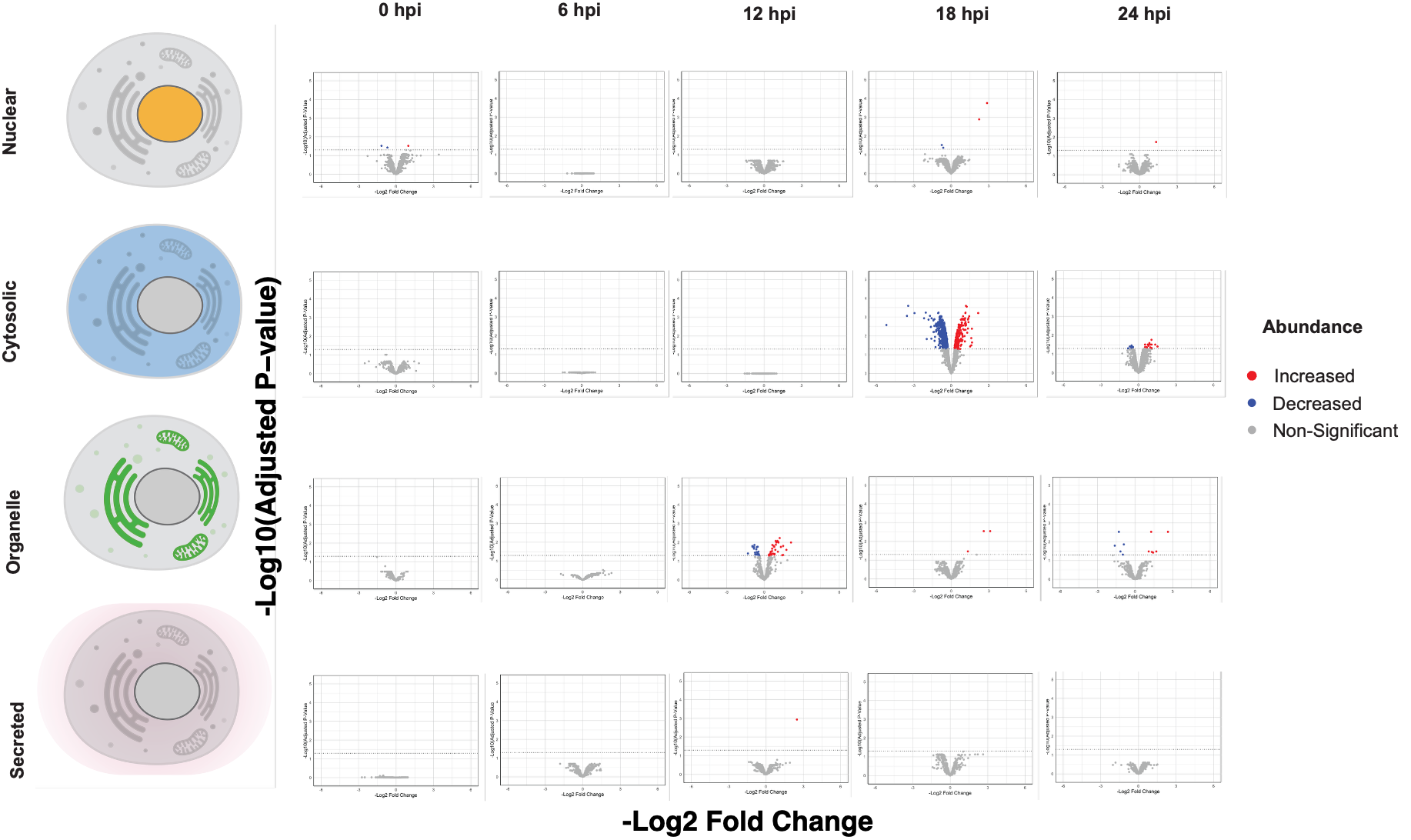

**Figure S1. Volcano plots of changes in protein abundance in subcellular fractions following PR8 infection compared to mock infection.** Individual volcano plots of differentially abundant proteins are shown for each timepoint (0, 6, 12, 18 and 24 h) post PR8 (H1N1) infection in each nuclear, cytosolic, organelle, or secreted fraction. Statistical comparisons were performed using the R package MSstatsTMT. Proteins significantly (Adjusted p-value ≤ 0.05) increased in abundance, red; proteins significantly decreased in abundance, blue; and proteins not significantly different, grey.

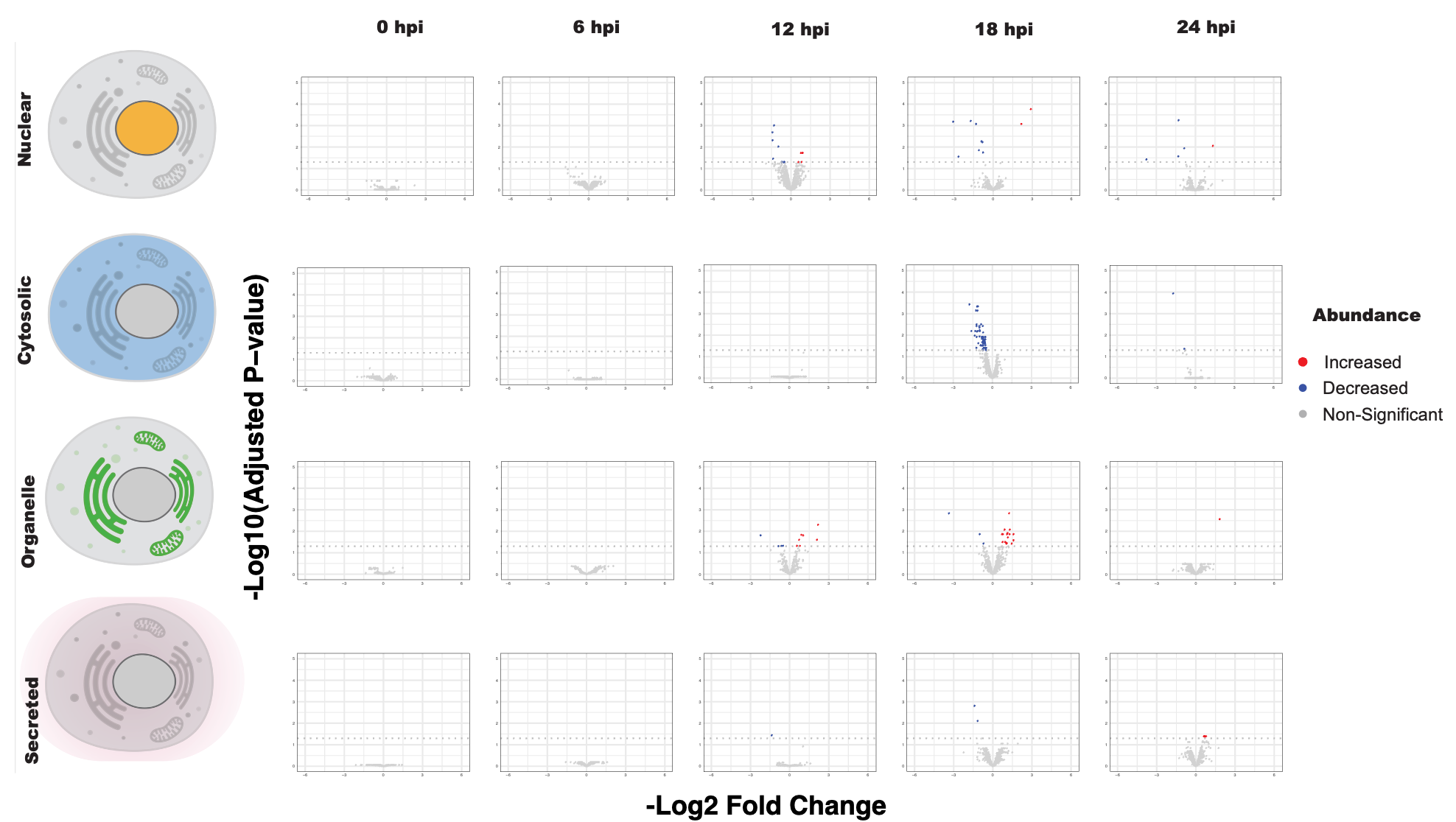

**Figure S2. Volcano plots of changes in protein abundance in subcellular fractions following PR8 infection compared to x31 infection.** Individual volcano plots of differentially abundant proteins are shown for each timepoint (0, 6, 12, 18 and 24 h) post PR8 (H1N1) infection as compared to x31 (H3N2) infection in each nuclear, cytosolic, organelle, or secreted fraction. Statistical comparisons were performed using MSstatsTMT. Proteins significantly (adjusted p < 0.05) increased in abundance, red; proteins significantly decreased in abundance, blue; and proteins not significantly different, grey.

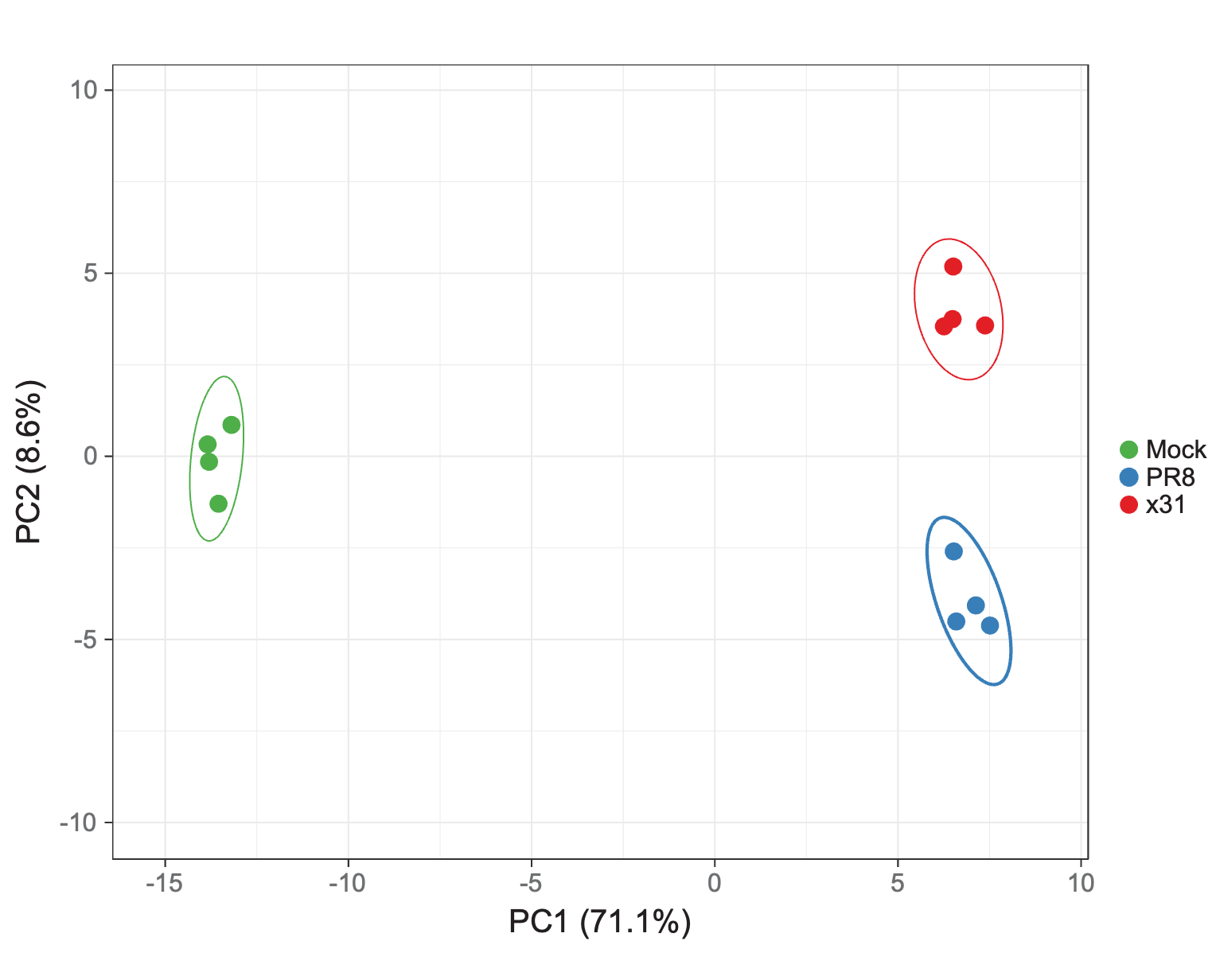
**Figure S3. Principal Component Analysis of the secreted *N*-glycome in IAV infection.** Principal component analysis of secreted *N-*glycan normalised abundance of A549 cells 24 hpi with PR8 (H1N1), in blue; x31 (H3N2), in red; and mock infection with naïve allantoic fluid in PBS, in green. *N*-glycans released from secreted proteins were separated and quantified using PGC-LC-MS/MS and were normalised to total glycan abundance.

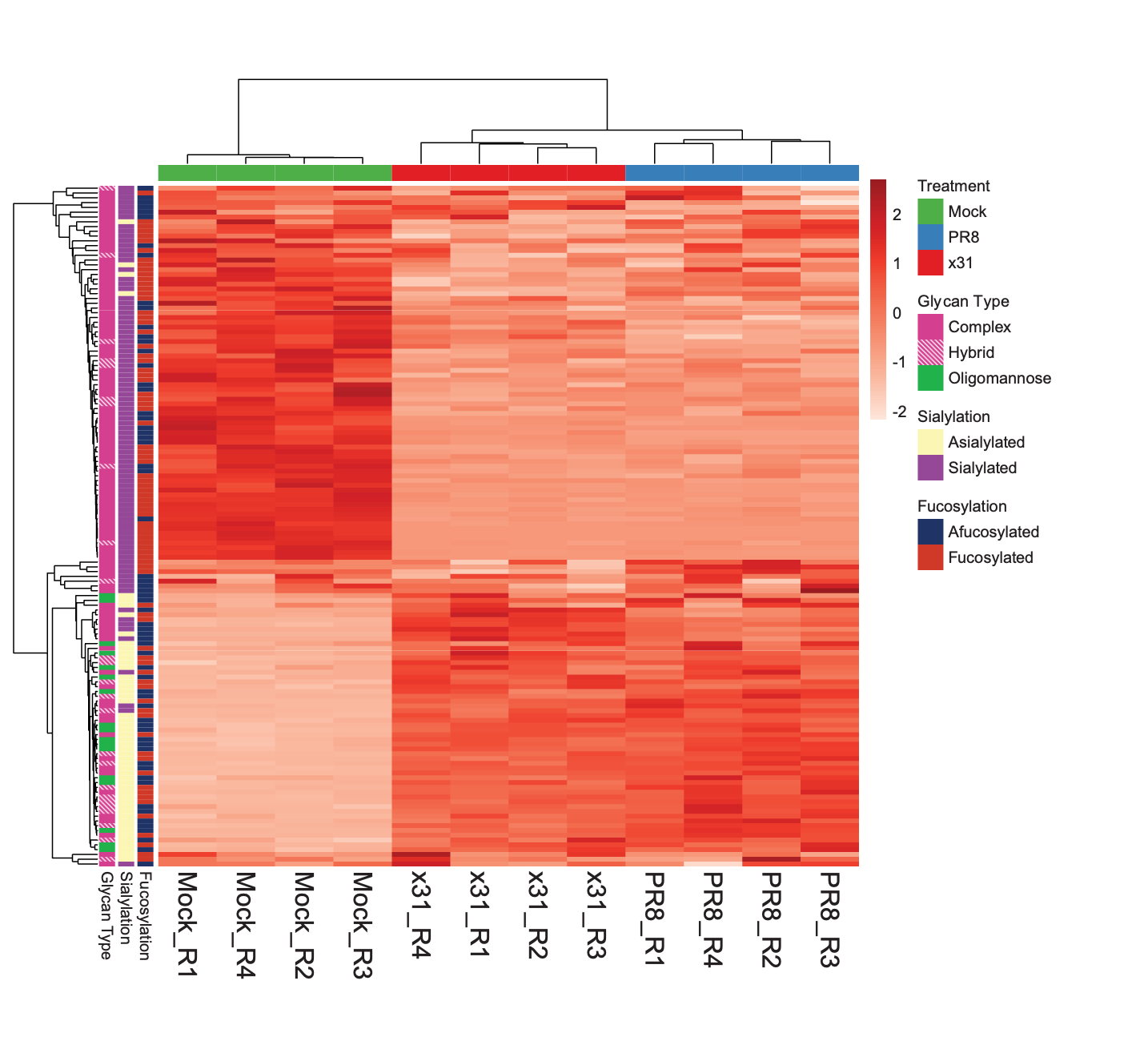
**Figure S4. Clustered heatmap of the secreted *N*-glycome in IAV infection.** Clustered heatmap of secreted *N*-glycans of A549 cells 24 hpi with PR8 (H1N1), in blue; x31 (H3N2), in red; and mock infection with naïve allantoic fluid in PBS, in green, coloured by z score of glycan normalised abundance. *N*-glycans released from secreted proteins were separated and quantified using PGC-LC-MS/MS and were normalised to total glycan abundance. Released glycan structures are stratified by their glycan class, sialylation and fucosylation status.

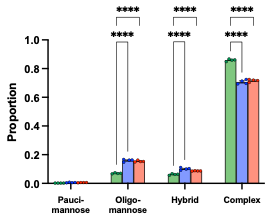
**Figure S5. *N*-glycan class abundance in the secreted *N*-glycome in IAV Infection.** Summed proportion of the major glycan classes of secreted *N*-glycans in A549 cells 24 hpi with PR8 (H1N1), in blue; x31 (H3N2), in red; and mock infection with naïve allantoic fluid in PBS, in green. *N*-glycans released from secreted proteins were separated and quantified using PGC-LC-MS/MS and were normalised to total glycan abundance. Values show the mean, errors bars the SEM. *, P ≤ 0.05; **, P ≤ 0.01; ***, P ≤ 0.001; ****, P ≤ 0.0001.

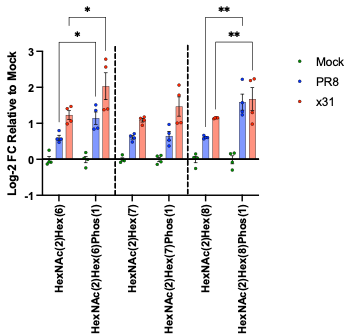
**Figure S6. Abundance of phosphorylated oligomannose *N*-glycans in the organelle fraction in IAV infection.** Log2FC relative to normalised abundance in mock-infected samples of oligomannose glycans with and without additional phosphorylation. Glycans are released from organelle proteins harvested from A549 cells 24 hpi with PR8 (H1N1), in blue; x31 (H3N2), in red; or mock infected with naïve allantoic fluid in PBS, in green. Values show the mean, error bars the SEM. *, P ≤ 0.05; **, P ≤ 0.01; ***, P ≤ 0.001; ****, P ≤ 0.0001.

**Figure S7. Glycan mass and extent of desialylation in secreted *N-*glycans in IAV infection. (A)** Correlation between the decrease in abundance (-Log2 FC) relative to mock of mono-, di- and tri-sialylated secreted *N-*glycans and their mass (Da) 24 hpi with PR8 (H1N1), in blue; x31 (H3N2), in red; and mock infection with naïve allantoic fluid in PBS, in green. *N*-glycans released from secreted proteins were separated and quantified using PGC-LC-MS/MS and were normalised to total glycan abundance. **(B)** Log2 FC in abundance relative to mock when summing glycans by HexNAc and Hex compositions (regardless of terminal sialylation or fucosylation). Values show mean, error bars SEM. *, P ≤ 0.05; **, P ≤ 0.01; ***, P ≤ 0.001; ****, P ≤ 0.0001.

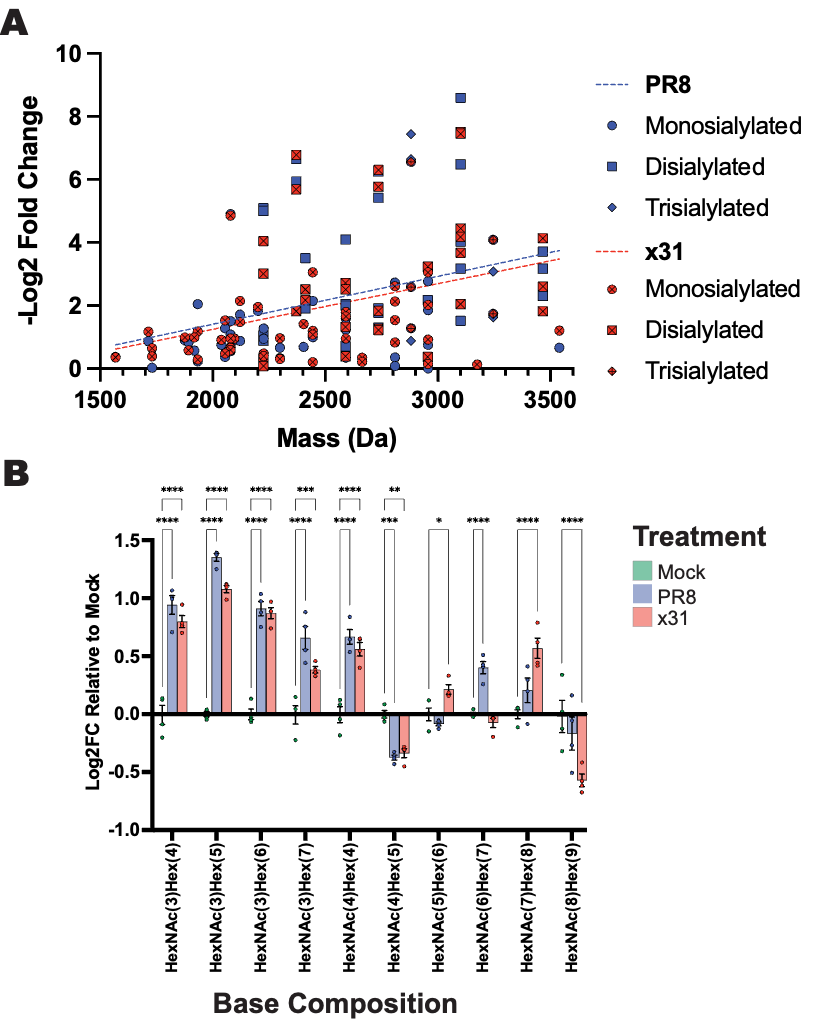

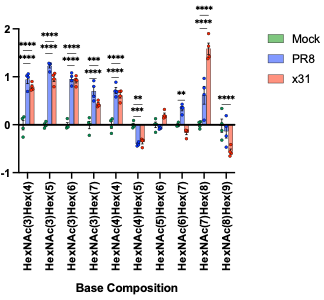

**Figure S8. Subcellular glycoproteomics in PR8 Infection**. **(A)** Volcano plots comparing Log2 FC and p-value of differentially abundant *N-*glycopeptides in organelle and secreted protein fractions every 6 h from 0 to 24 hpi with PR8 (H1N1) infection. *N*-glycopeptide abundances are normalised to their site-specific proportions.
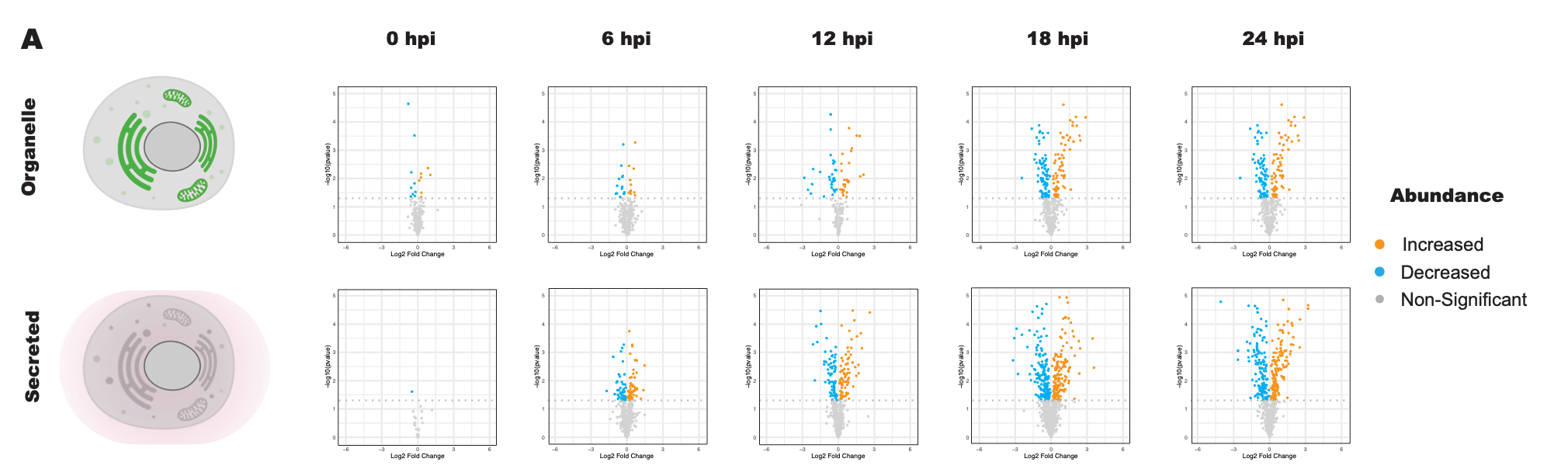

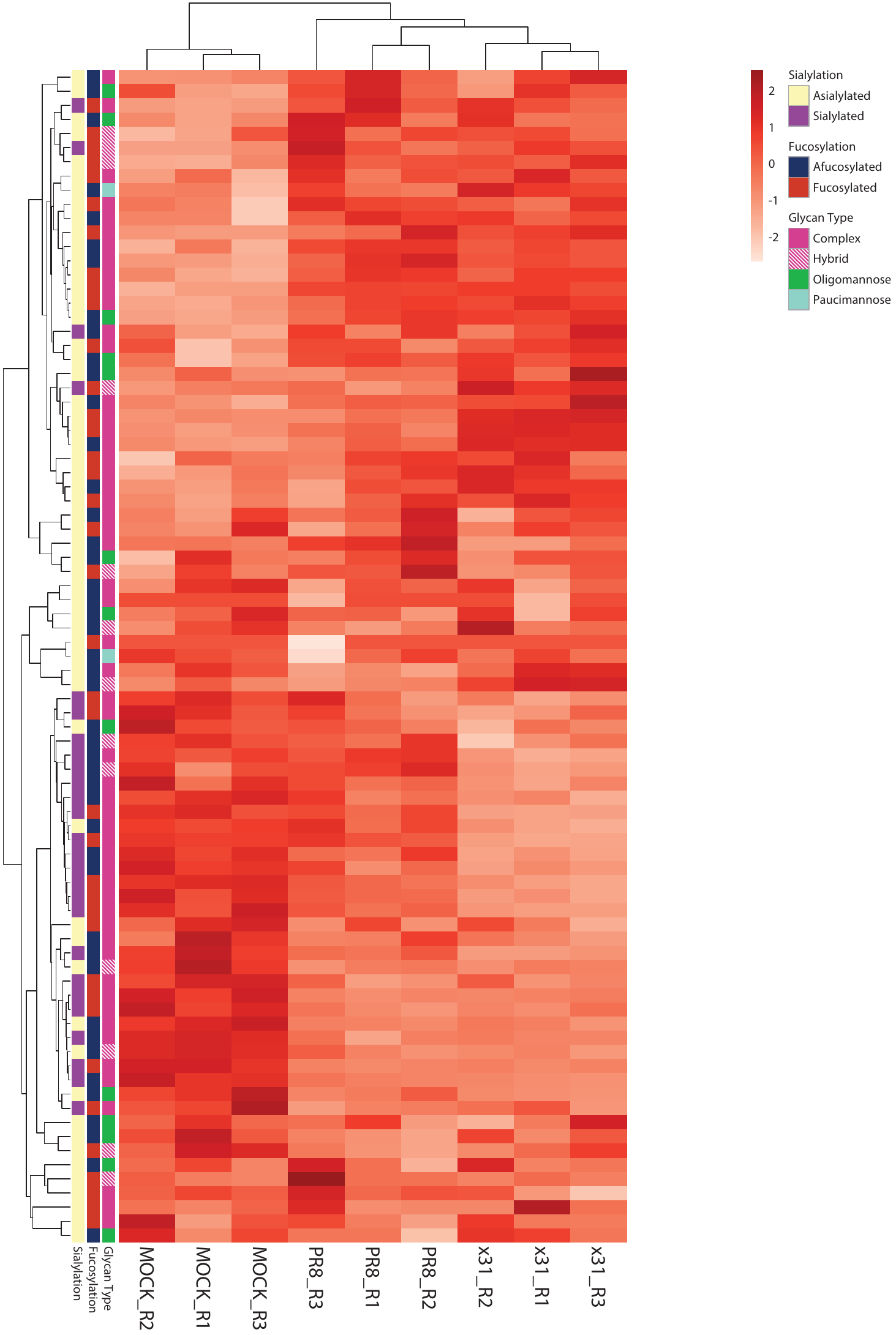
**Figure S9. Clustered heatmap of site-specific organelle glycopeptide abundances 12 h post IAV infection.** Clustered heatmap of the normalised abundance of organelle *N*-glycopeptides stratified by their sialylation status 12 hpi with PR8 (H1N1), in blue; x31 (H3N2), in red; and mock infection with naïve allantoic fluid in PBS, in green, coloured by z score of site normalised glycopeptide abundance.

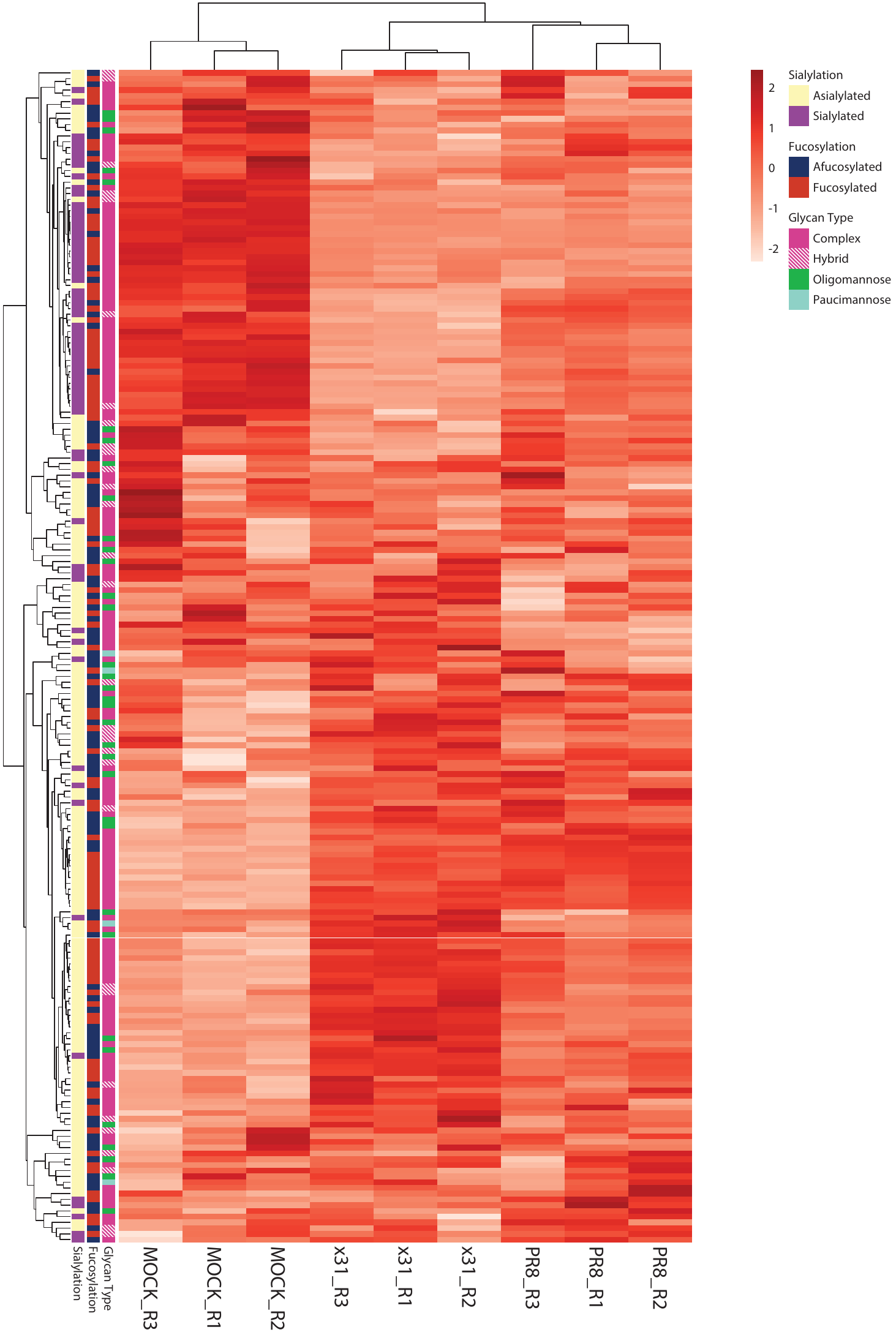
**Figure S10. Clustered heatmap of site-specific organelle glycopeptide abundances 18 h post IAV infection.** Clustered heatmap of the normalised abundance of organelle *N*-glycopeptides stratified by their sialylation status 18 hpi with PR8 (H1N1), in blue; x31 (H3N2), in red; and mock infection with naïve allantoic fluid in PBS, in green, coloured by z score of site normalised glycopeptide abundance.

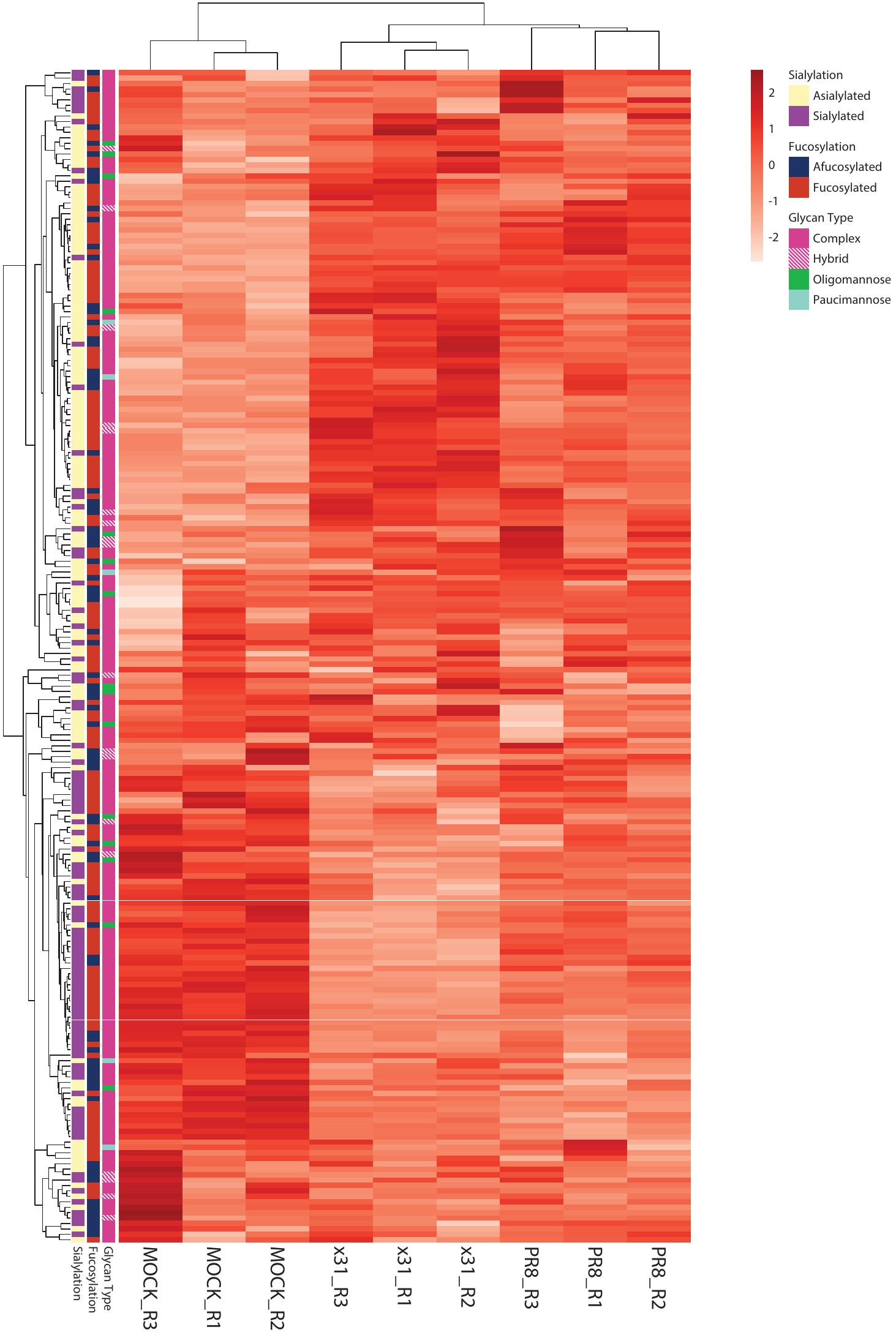
**Figure S11. Clustered heatmap of site-specific secreted glycopeptide abundances 12 h post IAV infection.** Clustered heatmap of the normalised abundance of secreted *N*-glycopeptides stratified by their sialylation status 12 hpi with PR8 (H1N1), in blue; x31 (H3N2), in red; and mock infection with naïve allantoic fluid in PBS, in green, coloured by z score of site normalised glycopeptide abundance.

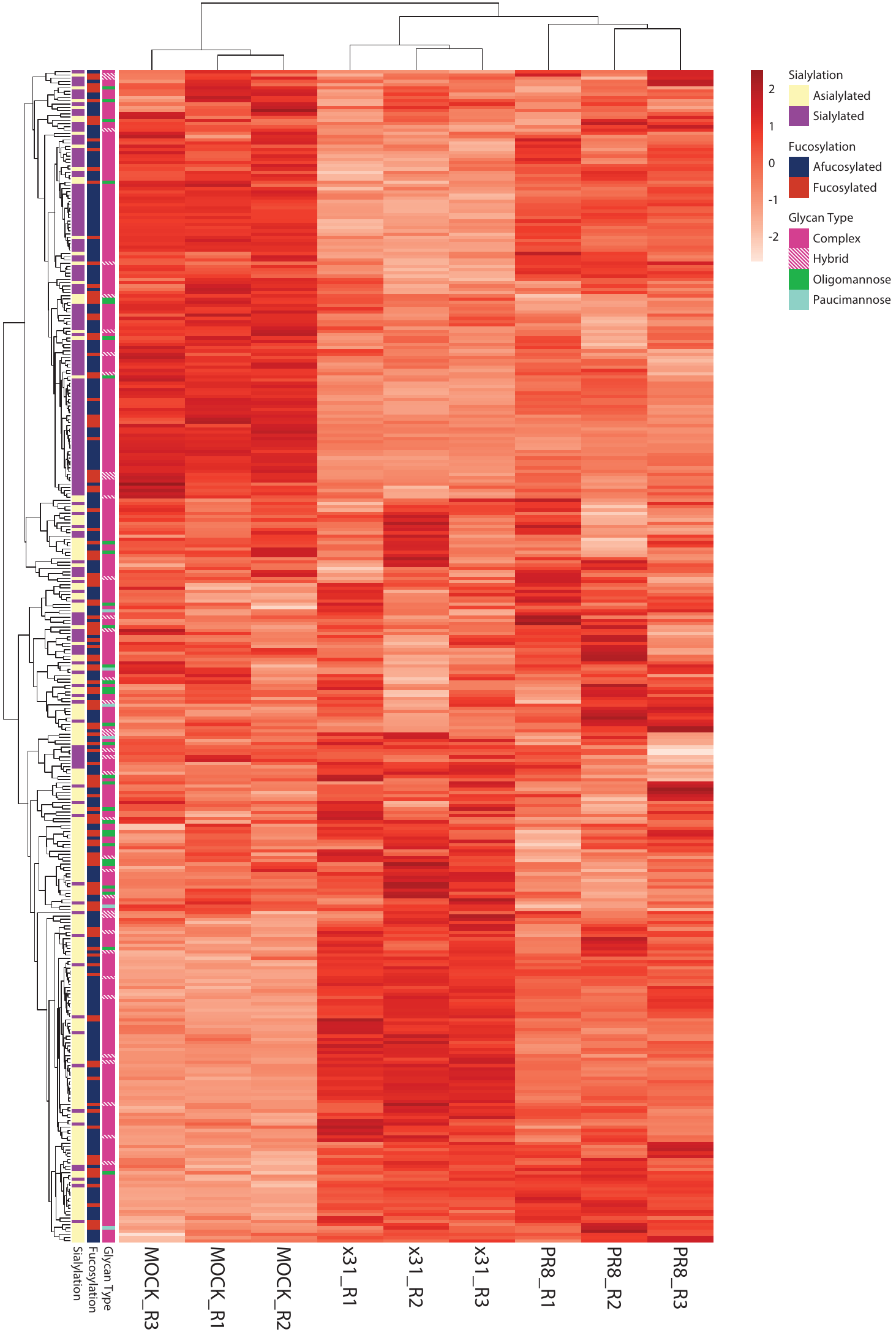
**Figure S12. Clustered heatmap of site-specific secreted glycopeptide abundances 18 h post IAV infection.** Clustered heatmap of the normalised abundance of secreted *N*-glycopeptides stratified by their sialylation status 18 hpi with PR8 (H1N1), in blue; x31 (H3N2), in red; and mock infection with naïve allantoic fluid in PBS, in green, coloured by z score of site normalised glycopeptide abundance.

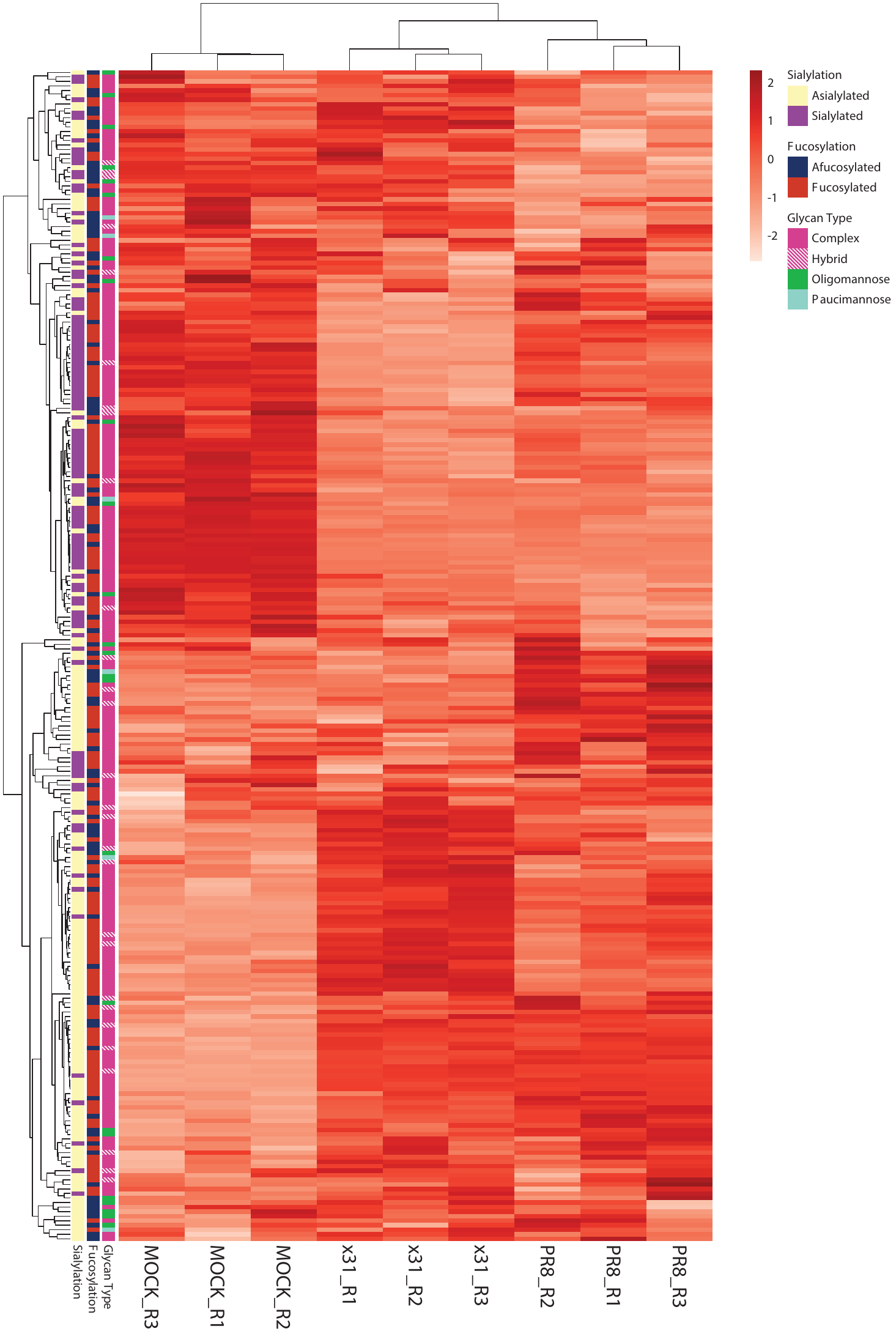
**Figure S13. Clustered heatmap of site-specific secreted glycopeptide abundances 24 h post IAV infection.** Clustered heatmap of the normalised abundance of secreted *N*-glycopeptides stratified by their sialylation status 24 hpi with PR8 (H1N1), in blue; x31 (H3N2), in red; and mock infection with naïve allantoic fluid in PBS, in green, coloured by z score of site normalised glycopeptide abundance.

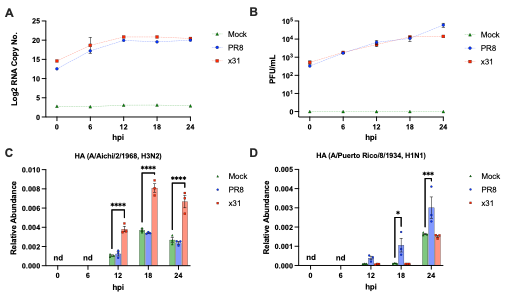
**Figure S14. Replication kinetics of influenza A virus infection. (A)** RNA copy number (Log2 transformed) of the A/Puerto Rico/8/1934 matrix gene determined using qRT-PCR analysis using the standard curve method and serial dilution of a matrix gene plasmid of known concentration on A549 cells 0, 6, 12, 18 and 24 hpi with PR8 (H1N1), in blue; and x31 (H3N2), in red; as well as mock infection with naïve allantoic fluid diluted in PBS, in green. **(B)** Viral infectious particle titres in plaque forming units/mL determined from plaque-forming assays of MDCK cells treated with serial dilutions of spent cell culture media from infections. Cells were cultured in a semi-solid agarose overlay that restricts viral spread to nearby cells only, resulting in the formation of visible and countable plaques with cell death. **(C-D)** Relative protein abundance is displayed for the hemagglutinin (HA) proteins of x31 **(C)** and PR8 **(D)**. Protein abundances were determined through liquid chromatography tandem mass spectrometry analysis and were normalised to the total protein abundance of human proteins identified within the sample. The use of ‘nd’ indicates time points where the HA protein was not detected. Values show the mean, error bars the SEM. *, P ≤ 0.05; **, P ≤ 0.01; ***, P ≤ 0.001; ****, P ≤ 0.0001.

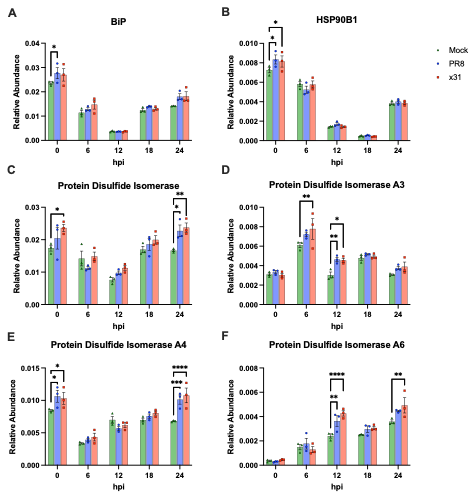
**Figure S15. Relative abundance of select Unfolded Protein Response-associated proteins in IAV infection.** Relative protein abundance of UPR associated proteins **(A)** BiP (P11021), **(B)** HSP90B1 (P14625), **(C)** protein disulfide isomerase (P07237), **(D)** protein disulfide isomerase A3 (P30101), **(E)** protein disulfide isomerase A4 (P13667) and **(F)** protein disulfide isomerase A6 (Q15084) in the organelle fraction of A549 cells at 0, 6, 12, 18 and 24 hpi with PR8 (H1N1), in blue; x31 (H3N2), in red; and mock infection with naïve allantoic fluid diluted in PBS, in green. Protein abundances were determined through liquid chromatography tandem mass spectrometry analysis and were normalised to the total protein abundance of human proteins identified within the sample. Values show the mean, error bars the SEM. *, P ≤ 0.05; **, P ≤ 0.01; ***, P ≤ 0.001; ****, P ≤ 0.0001.

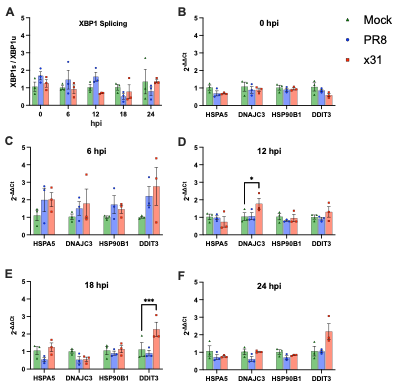
**Figure S16**. **Expression of Unfolded Protein Response-associated genes in IAV infection.** Fold change relative to mock of UPR associated genes of interest in A549 cells at 0, 6, 12, 18 and 24 hpi with PR8 (H1N1), in blue; x31 (H3N2), in red; and mock infection with naïve allantoic fluid diluted in PBS, in green. Fold change relative to mock is determined using qRT-PCR analysis using the ∆∆Ct method with GAPDH used as an endogenous control. **(A)** XBP1 splicing is determined by expression of spliced XBP1 mRNA (XBP1s) relative to the unspliced XBP1 mRNA (XBP1u). **(B-F)** Fold change in expression of the HSPA5, DNAJC3, HSP90B1 and DDIT3 genes at **(B)** 0 hpi, **(C)** 6 hpi, **(D)** 12 hpi, **(E)** 18 hpi and **(F)**24 hpi. Values show the mean, error bars the SEM. *, P ≤ 0.05; **, P ≤ 0.01; ***, P ≤ 0.001; ****, P ≤ 0.0001.
